# Deep Learning Dynamic Allostery of G-Protein-Coupled Receptors

**DOI:** 10.1101/2023.01.15.524128

**Authors:** Hung N. Do, Jinan Wang, Yinglong Miao

## Abstract

G-protein-coupled receptors (GPCRs) are the largest superfamily of human membrane proteins and represent primary targets of ∼1/3 of currently marketed drugs. Allosteric modulators have emerged as more selective drug candidates compared with orthosteric agonists and antagonists. However, many X-ray and cryo-EM structures of GPCRs resolved so far exhibit negligible differences upon binding of positive and negative allosteric modulators (PAMs and NAMs). Mechanism of dynamic allosteric modulation in GPCRs remains unclear. In this work, we have systematically mapped dynamic changes in free energy landscapes of GPCRs upon binding of allosteric modulators using the Gaussian accelerated molecular dynamics (GaMD), Deep Learning (DL) and free energy prOfiling Workflow (GLOW). A total of 18 available high-resolution experimental structures of allosteric modulator-bound class A and B GPCRs were collected for simulations. A number of 8 computational models were generated to examine selectivity of the modulators by changing their target receptors to different subtypes. All-atom GaMD simulations were performed for a total of 66 µs on 44 GPCR systems in the presence/absence of the modulator. DL and free energy calculations revealed significantly reduced conformational space of GPCRs upon modulator binding. While the modulator-free GPCRs often sampled multiple low-energy conformational states, the NAMs and PAMs confined the inactive and active agonist-G protein-bound GPCRs, respectively, to mostly only one specific conformation for signaling. Such cooperative effects were significantly reduced for binding of the selective modulators to “non-cognate” receptor subtypes in the computational models. Therefore, comprehensive DL of extensive GaMD simulations has revealed a general dynamic mechanism of GPCR allostery, which will greatly facilitate rational design of selective allosteric drugs of GPCRs.

## Introduction

G-protein-coupled receptors (GPCRs) are the largest superfamily of human membrane proteins with >800 members. GPCRs play key roles in cellular signaling and mediate various physiological activities, including vision, olfaction, taste, neurotransmission, endocrine, and immune responses^1^. They represent primary targets of ∼1/3 of currently marketed drugs^2^. GPCRs can be classified into six different classes, including class A (Rhodopsin-like), B (secretin receptors), C (metabotropic glutamate receptors (mGluRs)), D (fungal mating pheromone receptors), E (cyclic AMP receptors), and class F (frizzled/TAS2 receptors)^3,4^. GPCRs share a characteristic structural fold of seven transmembrane (TM) helices (TM1-TM7) connected by three extracellular loops (ECL1-ECL3) and three intracellular loops (ICL1-ICL3). For decades, the primary endogenous agonist-binding (“orthosteric”) site has been targeted for drug design of GPCR agonists, antagonists and inverse agonists^5^. However, the orthosteric site is usually highly conserved in different subtypes of GPCRs. An orthosteric drug often binds and activates/deactivates multiple GPCRs simultaneously with poor selectivity, thereby causing toxic side effects^6^.

Alternatively, allosteric modulators have been discovered to bind topographically distant (“allosteric”) sites of GPCRs with advantages^7-12^. They are able to modulate the binding affinity and signaling of orthosteric ligands, including positive and negative allosteric modulators (PAMs and NAMs)^13^. The allosteric effect has been shown to depend on the orthosteric probe^14^, with a “ceiling level” determined by the magnitude and direction of cooperativity between the orthosteric and allosteric ligands. Because the allosteric site is usually more divergent in residue sequences and conformations, allosteric modulators offer higher receptor selectivity than the orthosteric ligands. They serve as important chemical probes and promising selective therapeutics of GPCRs.

Important insights have been obtained using X-ray crystallography and cryo-electron microscopy (cryo-EM) about structural changes induced by allosteric modulator binding in certain GPCRs^7,9,10,15^. For class A GPCRs, binding of the LY2119620 PAM in the M_2_ muscarinic receptor (M_2_R) led to sidechain rotation of residue W7.35 and slight contraction of the receptor extracellular pocket, which was pre-formed in the active agonist-bound structure^16,17^. GPCR residues are numbered according to the Ballesteros-Weinstein scheme^18^. Binding of the muscarinic toxin MT7 NAM to the antagonist bound M_1_ muscarinic receptor (M_1_R) resulted in conformational changes in the ECL2, TM1, TM2, TM6 and TM7 extracellular domains, as well as the TM2 and TM6 intracellular domains^19^. In the free fatty acid receptor GPR40 (FFAR1), AgoPAM binding in a lipid-facing pocket formed by TM3-TM4-ICL2 induced conformational changes in the ICL2, TM4 and TM5 of the active receptor^20^. The ICL2 adopted a short helical conformation and the TM5 was shifted along its helical axis towards the extracellular side relative to the TM4^20^. A similar allosteric site was identified for binding of the NDT9513727 and Avacopan NAMs between TM3-TM4-TM5 on the lipid-exposed surface of the C5a_1_ receptor (C5AR1)^21^. For class B GPCRs, the LSN3160440 PAM was found to bind between the extracellular domains of TM1 and TM2 of the GLP-1 receptor (GLP1R)^22^. In the glucagon receptor (GLR), NAM binding outside of the 7TM bundle between TM6-TM7 restricted the outward movement of the TM6 intracellular domain required for activation and G-protein coupling of the receptor^23^. The ECL2 stretched to the central axis of the TM helical bundle, allowing for interactions from TM3 to TM6 and TM7 in the inactive class B GPCRs^23^. Despite remarkable advances, the X-ray and cryo-EM structures represent static snapshots of GPCRs. Many GPCR structures exhibit small/negligible differences in the absence and presence of allosteric modulators, notably for the A_1_AR^24^, M_4_R^25,26^, β_2_-adrenoceptor (β_2_AR)^27-29^, C5AR1^30^, CB_1_ cannabinoid receptor (CB_1_)^31^, chemokine receptor CCR2^32^, dopamine receptor 1 (D_1_R)^33^, GPBA receptor (GPBAR)^34^, and GLP1R^22,35,36^. A dynamic review has been suggested for allosteric modulation of GPCRs^37^. However, the dynamic mechanism of GPCR allostery remains unclear.

Molecular dynamics (MD) is a powerful computational technique for simulating biomolecular dynamics on an atomistic level^38^. For GPCRs, MD has been applied to simulate binding of both orthosteric and allosteric ligands^39-44^. Binding of known NAMs to the M_2_R was observed in conventional MD (cMD) simulations using the specialized supercomputer Anton^45^. The modulators formed cation-π interactions with aromatic residues in the receptor extracellular vestibule, which was confirmed by mutation experiments and later by X-ray structure of M_2_R recognized by a PAM^16^. Microsecond-timescale cMD simulations revealed mechanistic insights into allosteric modulation by Na^+^ in dopamine and opioid receptors^46,47^. Accelerated MD (aMD) simulations also captured Na^+^ binding to the highly conserved D2.50 allosteric site, which stabilized a muscarinic GPCR in the inactive state^48^. Recently, spontaneous binding of prototypical PAMs to the putative ECL2 allosteric site of the A_1_AR was captured in Gaussian accelerated molecular dynamics (GaMD) simulations^49^. Moreover, metadynamics simulations captured binding of the BMS-986187 PAM to the δ-opioid receptor^50^. Metadynamics and GaMD enhanced sampling simulations revealed positive binding cooperativity between allosteric and orthosteric ligands of the CCR2^51^. Despite these exciting advances, MD simulations of allosteric modulation have been limited to mostly few selected class A GPCRs^44^.

Recently, we have developed the GaMD, Deep Learning (DL) and free energy profiling workflow (GLOW) to predict molecular determinants and map free energy landscapes of biomolecules^52^. GaMD is an unconstrained enhanced sampling technique that works by applying a harmonic boost potential to smooth biomolecular potential energy surface^53^. Since this boost potential usually exhibits a near Gaussian distribution, cumulant expansion to the second order (“Gaussian approximation”) can be applied to achieve proper energy reweighting^54^. GaMD allows for simultaneous unconstrained enhanced sampling and free energy calculations of large biomolecules^53^. GaMD has been successfully demonstrated on enhanced sampling of ligand binding, protein folding, protein conformational changes, protein-membrane/peptide/protein/nucleic acid/carbohydrate interactions^55^. In GLOW, DL of image-transformed residue contact maps calculated from GaMD simulation frames allows us to identify important residue contacts by classic gradient-based pixel attribution in the saliency (attention) maps^52,56^. Finally, free energy profiles of these residue contacts are calculated through reweighting of GaMD simulations to characterize the biomolecular systems of interest^52^.

In this work, we have applied GLOW to systematically map dynamic changes in free energy landscapes of GPCRs upon binding of allosteric modulators (**Figure 1**). A total of 18 different high-resolution experimental structures of class A and B GPCRs are collected for modeling and 8 computational models were generated by changing target receptors of the modulators to different subtypes. Our comprehensive DL analysis of extensive GaMD simulations has provided important mechanistic insights into the dynamic allostery of GPCRs.

**Figure 1.**
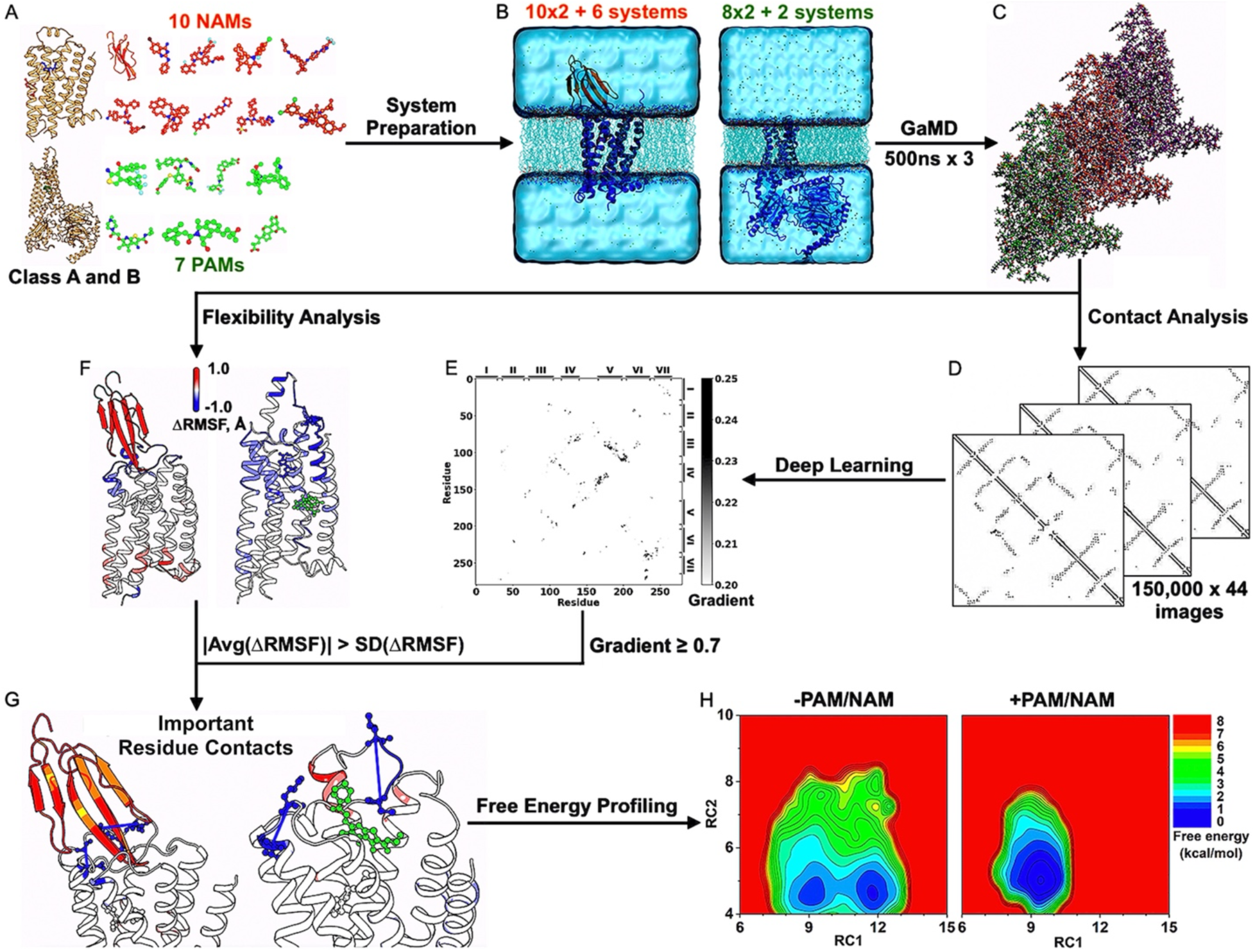
Workflow of deep learning dynamic allostery of GPCRs. Starting from 10 NAMs, 7 PAMs, 18 different experimental structures, and 8 computational models of class A and B GPCR-NAM/PAM complexes **(A)**, 2×10 structural + 6 model simulation systems of inactive antagonist-bound GPCRs in the presence/absence of NAM and 2×8 structural + 2 model simulation systems of active agonist-bound GPCRs in the presence/absence of PAM were built **(B)**. Three independent 500ns GaMD simulations were performed on each system **(C)**. Residue contact maps were calculated for 150,000 × 44 GaMD simulation frames **(D)** and analyzed by Deep Learning, yielding saliency (attention) maps of residue contact gradients **(E)**. Changes in root-mean-square fluctuations (ΔRMSFs) upon NAM/PAM binding in GPCRs were calculated from the GaMD simulations **(F)**. If the absolute average ΔRMSF calculated from three simulations of a residue was smaller than the standard deviation of ΔRMSF, the flexibility change for that residue was considered not significant and related residue pairs were neglected for further analysis. The characteristic residue contacts selected were those with ≥ 0.7 gradients and significant flexibility changes upon modulator binding **(G)**. They served as reaction coordinates for free energy profiling of dynamic allostery of GPCRs **(H)**.

## Methods

### Setup of GPCR simulation systems

A total of 18 unique experimental structures and 8 computational models of allosteric modulator-bound class A and B GPCRs were prepared for simulations (**Figure 1A** and **Supplementary Table 1**). The GPCR structures bound by NAMs included the MT7-bound M_1_R (PDB: 6WJC)^19^, Cmpd-15-bound β_2_AR (PDB: 5X7D)^27^, AS408-bound β_2_AR (PDB: 6OBA)^29^, NDT9513727-bound C5AR1 (PDB: 6C1Q)^30^, Avacopan-bound C5AR1 (PDB: 6C1R)^30^, ORG27569-bound CB_1_ receptor (PDB: 6KQI)^31^, GTPL9431-bound CCR2 (PDB: 5T1A)^32^, NNC0640-bound GLP1R (PDB: 5VEX)^35^, PF-06372222-bound GLP1R (PDB: 6LN2)^36^, and MK-0893-bound GLR (PDB: 5EE7)^23^. Six computational models of NAM-bound GPCRs included the MT7-bound M_2_R and M_4_R, which were built by aligning the 6WJC PDB structure of M_1_R to the 5ZK3^57^ and 5DSG^58^ PDB structures of M_2_R and M_4_R, respectively, and copying atomic coordinates of the atropine antagonist and MT7 NAM, as well as the Cmpd-15-bound α_1B_-adrenoceptor (α_1B_AR), α_2A_-adrenoceptor (α_2A_AR), α_2C_-adrenoceptor (α_2C_AR), and β_1_-adrenoceptor (β_1_AR), which were built by aligning the 5X7D PDB structure of β_2_AR to the 7B6W^59^, 6KUX^60^, 6KUW^61^, and 7BVQ^62^ PDB structures of α_1B_AR, α_2A_AR, α_2C_AR, and β_1_AR, respectively, and copying atomic coordinates of the carazolol antagonist and Cmpd-15 NAM. The GPCR structures bound by PAMs included the MIPS521-bound A_1_AR (PDB: 7LD3)^24^, LY2119620-bound M_2_R (PDB: 6OIK)^17^, LY2119620-bound M_4_R (PDB: 7V68)^25^, Cmpd-6FA-bound β_2_AR (PDB: 6N48)^28^, LY3154207-bound D_1_R (PDB: 7LJC)^33^, AgoPAM-bound FFAR1 (PDB: 5TZY)^20^, INT777-bound GPBAR (PDB: 7CFN)^34^, and LSN3160440-bound GLP1R (PDB: 6VCB)^22^. Two computational models of PAM-bound GPCRs included LY2119620-bound M_1_R, which was built by aligning the 6OIK PDB structure of M_2_R to the 6OIJ PDB structure of M_1_R^17^ and copying atomic coordinates of the LY2119620 PAM, and LY32154207-bound D_2_ receptor (D_2_R), which was built by aligning the 7LJC PDB structure of D_1_R to the 7JVR PDB structure of D_2_R^63^ and copying atomic coordinates of the SKF-81297 agonist and LY3154207 PAM.

SWISS-MODEL^64^ homology modeling was applied to restore missing residues in the GPCR structures and models, particularly in the ECL2, ICL2, and ECL3. Charges of the ligands were listed in **Supplementary Table 1**. All water and heteroatom molecules except the ligands and receptor-bound ions (including the sodium ion in the 6C1R PDB structure of C5AR1 and zinc ion in the 6LN2 PDB structure of GLP-1 receptor) were removed from the structures. The GPCR complexes were embedded in POPC membrane lipid bilayers and solvated in 0.15M NaCl (**Figure 1B**). The AMBER^65^ force field parameter sets were used for our GaMD simulations, specifically ff19SB^66^ for proteins, GAFF2^67^ for ligands, LIPID17 for lipids, and TIP3P^68^ for water, except for the A_1_AR simulations as obtained from a previous study^24^ where the CHARMM36m^69^ force field parameter set was used.

### Simulation protocols

All-atom dual-boost GaMD simulations^53^ were performed on the GPCR structures and models with and without allosteric modulators (**Figure 1C** and **Supplementary Table 1**). The simulations of the A_1_AR with and without the MIPS521 PAM were obtained from a previous study^24^. GaMD simulations of the other systems followed a similar protocol. The systems were energetically minimized for 5000 steps using the steepest-descent algorithm and equilibrated with the constant number, volume, and temperature (NVT) ensemble with 310 K. They were further equilibrated for 375 ps at 310 K with the constant number, pressure, and temperature (NPT) ensemble. The cMD simulations were then performed for 10 ns using the NPT ensemble with constant surface tension at 1 atm pressure and 310 K temperature. GaMD implemented in GPU version of AMBER 20^53,70,71^ was applied to simulate the GPCR systems. The simulations involved an initial short cMD of 4-10 ns to calculate GaMD acceleration parameters and GaMD equilibration of added boost potential for 16-40 ns. Three independent 500-ns GaMD production simulations with randomized initial atomic velocities were performed for each system at the “dual-boost” level, with one boost potential applied to the dihedral energetic term and the other to the total potential energetic term. The reference energy was set to the lower bound *E* = *V*_*max*_, and the upper limit of the boost potential standard deviation, **σ**_0_, was set to 6.0 kcal/mol for both the dihedral and total potential energetic terms. The GaMD simulations were summarized in **Supplementary Table 1**.

### Deep Learning and free energy profiling of GaMD simulations with the GLOW workflow

GLOW^52^ was applied to systematically analyze GPCR allostery (**Figure 1D-1E**). Residue contact maps of 6,600,000 GaMD simulation frames obtained from 66µs GaMD simulations were calculated and transformed into images for DL (**Figure 1D**). A contact definition of ≤4.5 Å between any heavy atoms in two protein residues was used. For DL, 80% of the residue contact map images were randomly assigned to the training set, while the remaining 20% were put in the validation set for each GPCR. The residue contact map images of each GPCR system were separated into four different classes for DL analysis based on the absence and presence of NAMs and PAMs, including NAM-free (“Antagonist”), NAM-bound (“AntagonistNAM”), PAM-free (“Agonist”), and PAM-bound (“AgonistPAM”). DL models of two-dimensional (2D) convolutional neural networks were built to classify the frame images with and without the allosteric modulator bound for each GPCR subfamily. Saliency (attention) maps of residue contact gradients were calculated through backpropagation by vanilla gradient-based pixel attribution^56^ using the residue contact map of the most populated structural cluster of each GPCR system (**Figure 1E**). The hierarchical agglomerative clustering algorithm was used to cluster snapshots of receptor conformations with all GaMD production simulations combined for each system^52^.

Furthermore, root-mean-square fluctuations (RMSFs) of the receptors and orthosteric ligands within the GPCR complexes were calculated by averaging the RMSFs calculated from individual GaMD simulations of each GPCR system. Changes in the RMSFs (ΔRMSF) upon binding of allosteric modulators were calculated by subtracting the RMSFs of GPCRs without modulators from those with modulators bound (**Figure 1F**). If the absolute average of ΔRMSF calculated from three simulations of a residue was smaller than the corresponding standard deviation, the flexibility change for that residue was considered not significant and related residue pairs were neglected for further analysis. Important residue contacts were selected with the highest contact gradients in the attention maps from DL and significant changes in the GPCR residue flexibility upon modulator binding (**Figures 1G-1H**). They were finally used as reaction coordinates (RCs) to calculate free energy profiles by reweighting the GaMD simulations using the *PyReweighting* toolkit^52-54^, with bin sizes of 0.5-1.0 Å and cutoff of 100-500 frames in one bin.

## Results

### GaMD simulations on effects of allosteric modulator binding to GPCRs

All-atom GaMD simulations were obtained for systems of the A_1_AR, M_1_R, M_2_R, M_4_R, α_1B_AR, α_2A_AR, α_2C_AR, β_1_AR, β_2_AR, C5AR1, CB_1_, CCR2, D_1_R, D_2_R, FFAR1, GPBAR, GLP1R, and GLR. GaMD simulations performed in this study recorded overall similar averages (∼13-16 kcal/mol) and standard deviations (∼4-5 kcal/mol) of boost potentials across the different GPCR systems, except for the A_1_AR simulations that were obtained from a previous study^24^ using a different force field parameter set^69^ (**Supplementary Table 1**). We first examined structural dynamics of the GPCR orthosteric and allosteric ligands. Time courses of the orthosteric and allosteric ligand RMSDs relative to the simulation starting structures were plotted in **Supplementary Figures 1-3**. In most of the GPCR systems, the orthosteric ligands underwent similar structural deviations in the absence and presence of allosteric modulators during the GaMD simulations (**Supplementary Figures 1-3**). This was consistent with previous findings that modulator binding mostly does not cause large changes in the X-ray and cryo-EM structures of the GPCRs^22,24-36^. However, a number of orthosteric ligands, including adenosine in A_1_AR and PE5 in GLR exhibited significantly smaller structural deviations in the presence of the MIPS521 PAM and MK-0893 NAM, respectively (**Supplementary Figures 2A** and **3D**).

**Figure 2.**
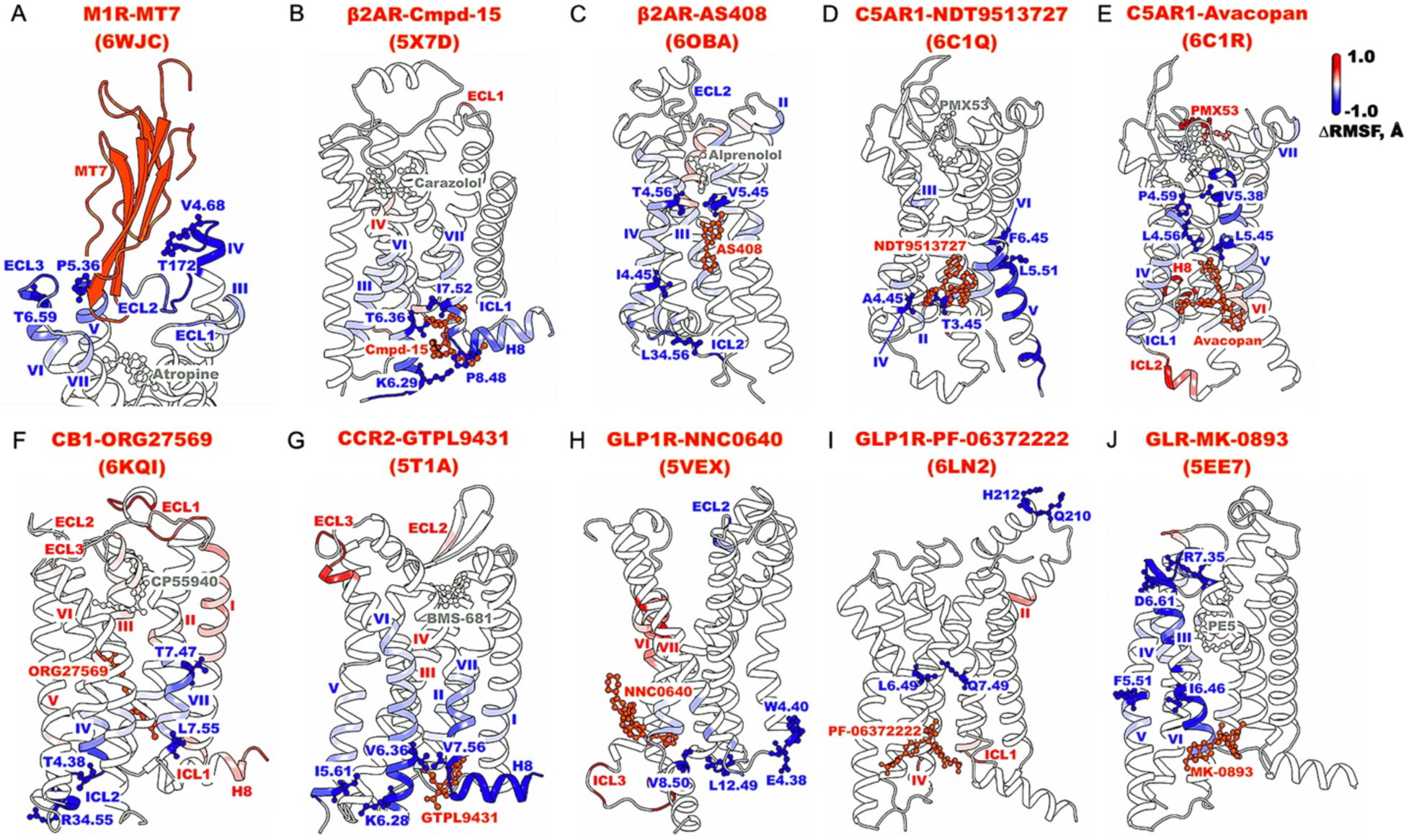
Characteristic residue contacts in the negative allosteric modulation of class A and B GPCRs calculated from GaMD simulations of the MT7-bound M_1_R (PDB: 6WJC) **(A)**, Cmpd-15-bound β_2_AR (PDB: 5X7D) **(B)**, AS408-bound β_2_AR (PDB: 6OBA) **(C)**, NDT9513727-bound C5AR1 (PDB: 6C1Q) **(D)**, Avacopan-bound C5AR1 (PDB: 6C1R) **(E)**, ORG27569-bound CB_1_ (PDB: 6KQI) **(F)**, GTPL9431-bound CCR2 (PDB: 5T1A) **(G)**, NNC0640-bound GLP1R (PDB: 5VEX) **(H)**, PF-06372222-bound GLP1R (PDB: 6LN2) **(I)**, and MK-0893-bound GLR (PDB: 5EE7) **(J)**. The seven TM helices are labeled I-VII, H8 for helix 8, ECL1-ECL3 for extracellular loops 1-3, and ICL1-ICL3 for intracellular loops 1-3. A color scale of -1.0 (blue) to 0 (white) to 1.0 (red) is used to show the ΔRMSF upon NAM binding, and NAMs are colored orange.

In general, binding of allosteric modulators reduced fluctuations of the orthosteric ligands and GPCRs as shown in **Figures 2-3** and **Supplementary Figures 4-5**. NAM binding primarily stabilized the allosteric binding pockets, with additional reduced flexibility observed in the extracellular and/or intracellular domains of the receptors (**Figure 2** and **Supplementary Figure 4**). In particular, binding of MT7 significantly reduced fluctuations of the extracellular mouth between ECL2 and ECL3 in M_1_R (**Figure 2A**). Binding of Cmpd-15 in the β_2_AR and GTPL9431 in the CCR2 reduced fluctuations of the allosteric pocket formed by TM6, TM7, ICL4, and H8 (**Figure 2B, 2G**). Notably, AS408, NDT9513727, and Avacopan bound to a similar TM3-TM4-TM5 region on the lipid-facing surface of the β_2_AR and C5AR1. They reduced fluctuations of the intracellular domains of TM3, TM4, and TM5 (**Figure 2C-2E**), as well as ICL2 in β_2_AR (**Figure 2C**). This was consistent with recent finding that the NDT9513727 NAM stabilized TM5 through the hydrophobic stacking between TM4 and TM5^72^. Binding of NNC0640 to GLP1R and MK-0893 to GLR stabilized the lipid-facing pocket on the intracellular domains of TM7 and TM6, respectively (**Figure 2H, 2J** and **Supplementary Figure 4H, 4J**). While PF-06372222 also bound to a similar region in GLP1R, no significant flexibility change was observed in the receptor likely because the modulator-free receptor was already stable (**Figure 2I** and **Supplementary Figure 4I**). Lastly, binding of ORG27569 to CB_1_ reduced fluctuations of the TM4, TM6, TM7 intracellular domains and ICL2 (**Figure 2F**).

Binding of PAMs to GPCRs generally reduced fluctuations of the extracellular domains, orthosteric agonist-binding pocket, and/or intracellular G-protein coupling domains (**Figure 3** and **Supplementary Figure 5**). Specifically, binding of MIPS521 to the A_1_AR significantly reduced fluctuations of the adenosine agonist, the orthosteric pocket constituted by extracellular domains of TM2, ECL1, ECL2, TM3, TM5, TM6, ECL3, and TM7, and the intracellular domains of TM5, TM7, and ICL2 (**Figure 3A** and **Supplementary Figure 5A**), which was consistent with the previous observation that the PAM stabilized the adenosine-A_1_AR-G-protein complex^24^. LY2119620 binding to the M_2_R reduced flexibility of TM2, TM7, ECL1-ECL3 and H8 (**Figure 3B** and **Supplementary Figure 5B**). In the M_4_R, LY2119620 binding significantly reduced flexibility of the G-protein coupling domains of ICL1, TM3, TM5, TM6, and H8, while stabilized ECL3 to a lesser extent (**Figure 3C** and **Supplementary Figure 5C**). In the β_2_AR, Cmpd-6FA binding reduced fluctuations of orthosteric residues in the ECL1 and TM7 extracellular end as well as the intracellular domains of ICL1, TM3, and ICL2 (**Figure 3D** and **Supplementary Figure 5D**). Binding of LY3154207 to the D_1_R reduced fluctuations of the TM1 extracellular domain and ECL3, as well as the TM6 intracellular domain and ICL2 (**Figure 3E** and **Supplementary Figure 5E**). Similar to the 6N48 PDB structure of β_2_AR, the 5TZY PDB structure of AgoPAM-bound FFAR1 did not have a G-protein bound. Binding of AgoPAM reduced flexibility of the ECL1, ECL2, ECL3 and TM1, TM2, TM3, and TM5 extracellular ends as well as the ICL2, and TM3 and TM4 intracellular ends, which constituted the G-protein binding pocket (**Figure 3F** and **Supplementary Figure 5F**). The stabilization of ICL2 observed in the AgoPAM-bound FFAR1 was in good agreement with the structural data of the PAM-bound 5TZY and PAM-free 5TZR PDB structures^20^. In particular, the ICL2 adopted a short helical conformation in the PAM-bound 5TZY PDB and was completely missing in the PAM-free 5TZR PDB structure of FFAR1^20^. Two molecules of INT777 with -1 charge served as the orthosteric and allosteric ligands of GPBAR, which potentially caused electrostatic repulsion and destabilized the orthosteric INT777 and most of the orthosteric residues (**Figure 3G** and **Supplementary Figure 5G**). In fact, Dror et al. uncovered that electrostatic repulsion between orthosteric and allosteric ligands could weaken the binding of one in the presence of the other^45^. Even so, binding of INT777 to the allosteric site reduced fluctuations of ECL2 and the TM4 extracellular end, as well as the TM5 and TM6 intracellular ends (**Figure 3G** and **Supplementary Figure 5G**). Finally, the LSN3160440 PAM binding significantly reduced fluctuations of the large orthosteric pocket of GLP1R formed by TM2, ECL1, ECL2, TM4, and ECL3, and the G-protein coupling domains in TM5 and TM6 (**Figure 3H** and **Supplementary Figure 5H**).

**Figure 3.**
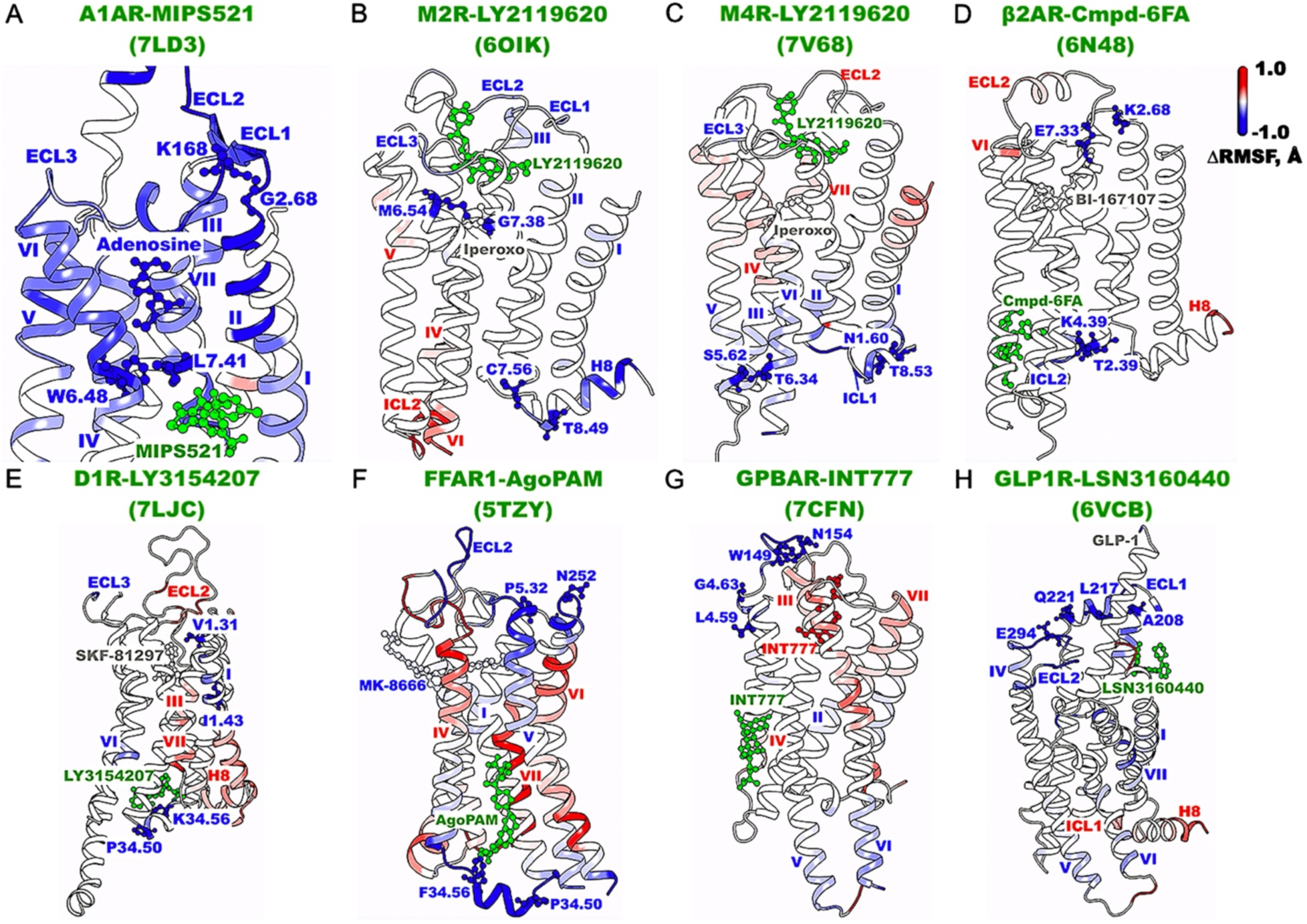
Characteristic residue contacts in the positive allosteric modulation of class A and B GPCRs calculated from GaMD simulations of. the MIPS521-bound A_1_AR (PDB: 7LD3) **(A)**, LY2119620-bound M_2_R (PDB: 6OIK) **(B)**, LY2119620-bound M_4_R (PDB: 7V68) **(C)**, Cmpd-6FA-bound β_2_AR (PDB: 6N48) **(D)**, LY3154207-bound D_1_R (PDB: 7LJC) **(E)**, AgoPAM-bound FFAR1 (PDB: 5TZY) **(F)**, INT777-bound GPBAR (PDB: 7CFN) **(G)**, and LSN3160440-bound GLP1R **(H)**. The seven TM helices are labeled I-VII, H8 for helix 8, ECL1-ECL3 for extracellular loops 1-3, and ICL1-ICL3 for intracellular loops 1-3. A color scale of -1.0 (blue) to 0 (white) to 1.0 (red) is used to show the ΔRMSF upon PAM binding, and PAMs are colored green.

**Figure 4.**
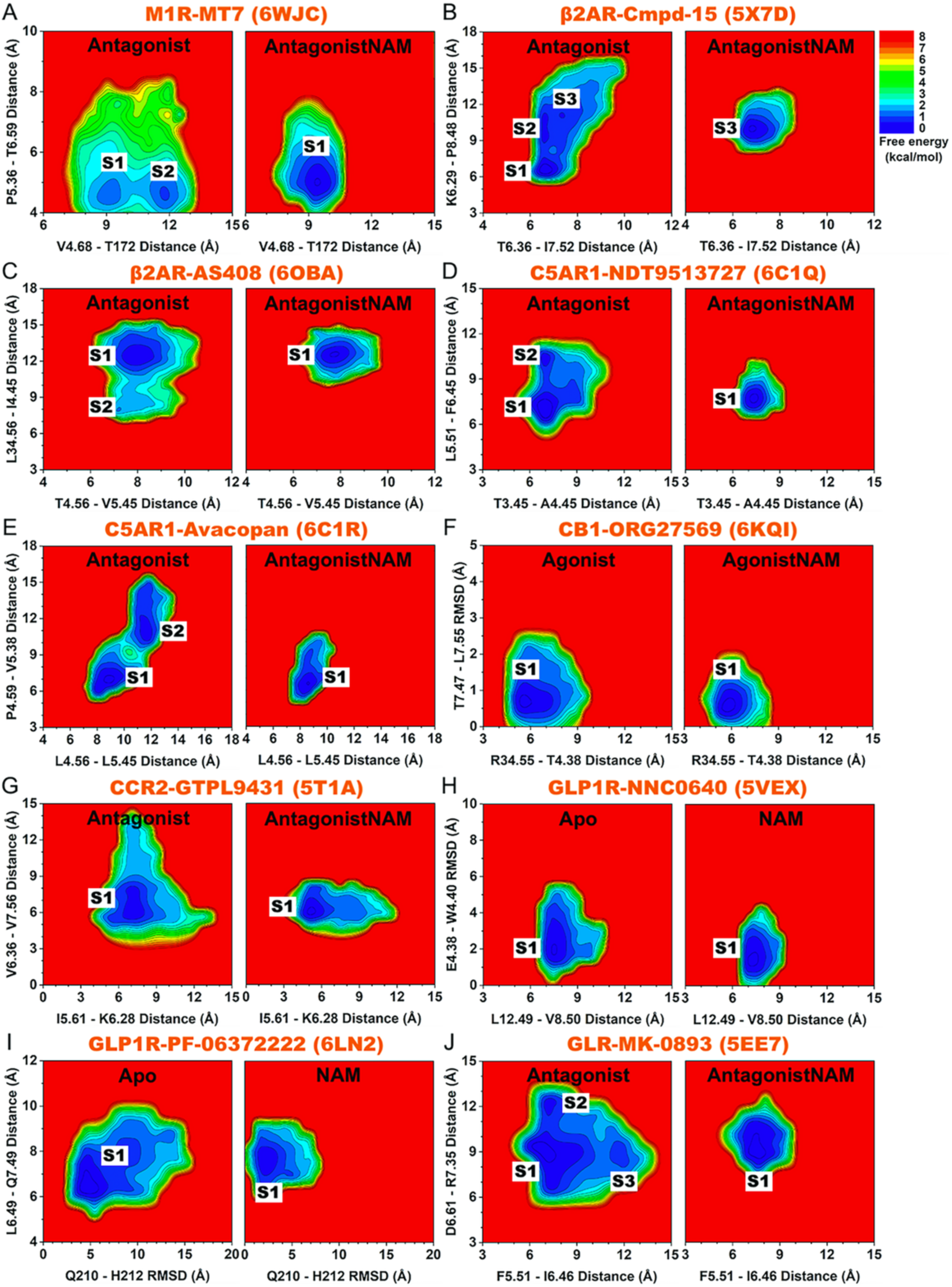
2D free energy profiles of characteristic residue contacts in the allosteric modulation of class A and B GPCRs bound by NAMs. **(A)** The C_α_-atom distances between V4.68-T172^ECL2^ and P5.36-T6.39 in the M_1_R without and with the MT7 NAM. The inactive antagonist-bound GPCR without and with NAM are denoted “Antagonist” and “AntagonistNAM”, respectively. **(B)** The C_α_-atom distances between T6.36-I7.52 and K6.29-P8.48 in the β_2_AR without and with the Cmpd-15 NAM. **(C)** The C_α_-atom distances between T4.56-V5.45 and L34.56^ICL2^-I4.45 in the β_2_AR without and with the AS408 NAM. **(D)** The C_α_-atom distances between T3.45-A4.45 and L5.51-F6.45 in the C5AR1 without and with the NDT9513727 NAM. **(E)** The C_α_-atom distances between L4.56-L5.45 and P4.59-V5.38 in the C5AR1 without and with the Avacopan NAM. **(F)** The C_α_-atom distance between R34.55^ICL2^-T4.38 and RMSD of T7.47-L7.55 relative to the 6KQI PDB structure in the CB_1_ receptor without and with the ORG27569 NAM. The agonist-bound GPCR without and with NAM are denoted “Agonist” and “AgonistNAM”, respectively. **(I)** The C_α_-atom distances between I5.61-K6.28 and V6.36-V7.56 in the CCR2 without and with the GTPL9431 NAM. **(J)** The C_α_-atom distance between L12.49^ICL1^-V8.50 and RMSD of E4.38-W4.40 relative to the 5VEX PDB structure in the GLP1R without and with the NNC0640 NAM. The apo GPCR without and with NAM are denoted “Apo” and “NAM”, respectively. **(K)** The RMSD of Q210^ECL1^-H212^ECL1^ relative to the 6LN2 PDB structure and C_α_-atom distance between L6.49-Q7.49 in the GLP1R without and with the PF-06372222 NAM. **(L)** The C_α_-atom distances between F5.51-I6.46 and D6.61-R7.35 in the GLR without and with the MK-0893 NAM.

**Figure 5.**
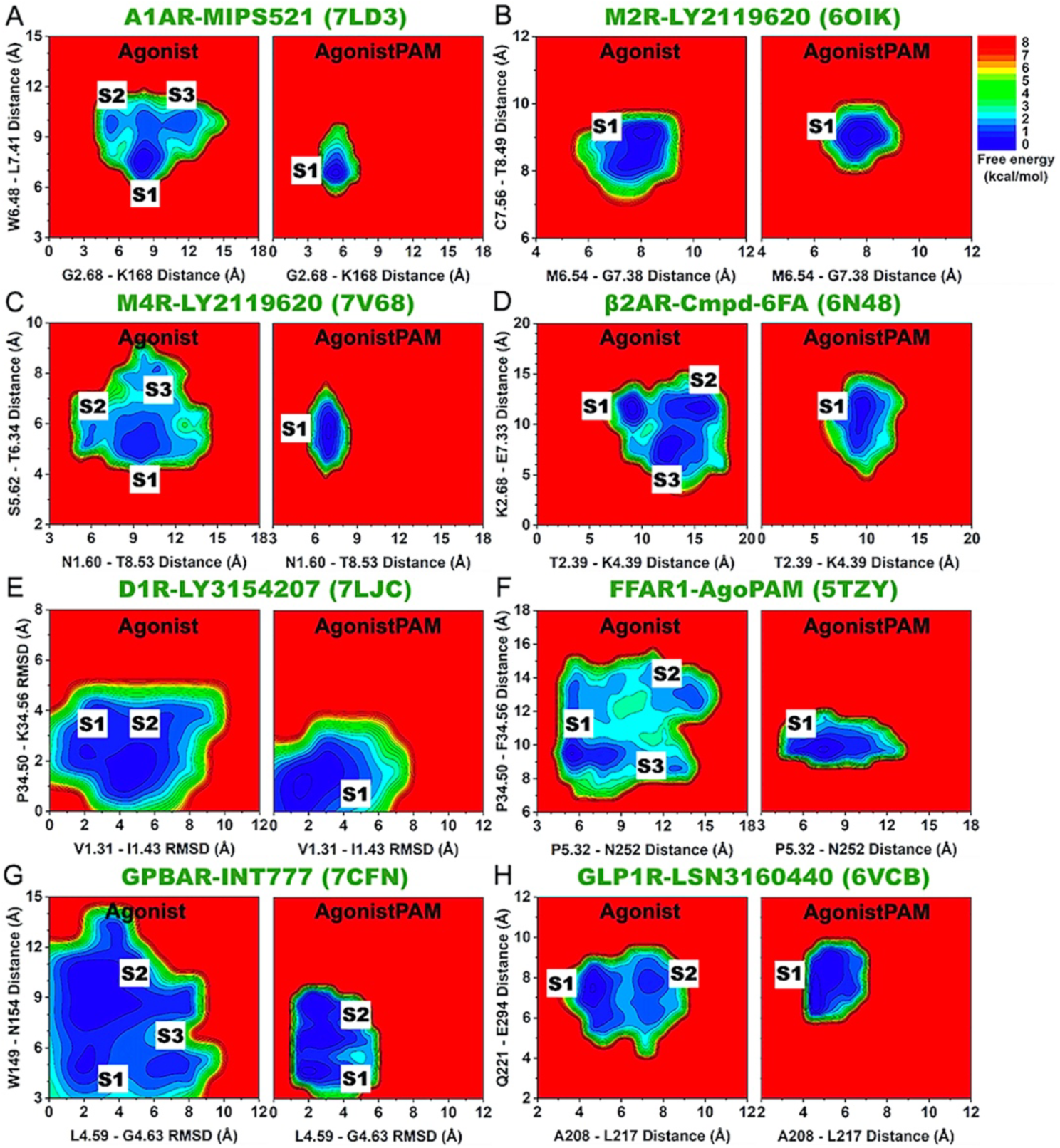
2D free energy profiles of characteristic residues in the allosteric modulation of class A and B GPCRs bound by PAMs. **(A)** The C_α_-atom distances between G2.68-K168^ECL2^ and W6.48-L7.41 in the A_1_AR without and with the MIPS521 PAM. The active agonist-bound GPCR without and with PAM are denoted “Agonist” and “AgonistPAM”, respectively. **(B)** The C_α_-atom distances between M6.54-G7.38 and C7.56-T8.49 in the M_2_R without and with the LY2119620 PAM. **(C)** The C_α_-atom distances between N1.60-T8.53 and S5.62-T6.34 in the M_4_R without and with the LY2119620 PAM. **(D)** The C_α_-atom distances between T2.39-K4.39 and K2.68-E7.33 in the β_2_AR without and with the Cmpd-6FA PAM. **(E)** The RMSDs of V1.31-I1.43 and P34.50^ICL2^-K34.56^ICL2^ relative to the 7LJC PDB structure in the D_1_R without and with the LY3154207 PAM. **(F)** The C_α_-atom distances between P5.32-N252^ECL3^ and P34.50^ICL2^-F34.56^ICL2^ in the FFAR1 without and with the AgoPAM PAM. **(G)** The RMSD of L4.59-G4.63 and C_α_-atom distance between W149^ECL2^-N154^ECL2^ in the GPBAR without and with the INT777 PAM. **(H)** The C_α_-atom distances between A208^ECL1^-L217^ECL1^ and Q221^ECL1^-E294^ECL2^ in the GLP1R without and with the LSN3160440 PAM.

### Deep Learning important residue contacts underlying allosteric modulation of GPCRs

DL models were built from image-transformed residue contact maps calculated from GaMD simulations of the modulator-free and modulator-bound systems for each GPCR subfamily. Classification of GPCRs bound by “Antagonist”, “AntagonistNAM”, “Agonist”, “AgonistPAM” was carried out with high accuracies on both the training and validation sets after 15 epochs for all GPCRs (**Supplementary Figures 6** and **7**). The saliency (attention) residue contact maps of gradients were calculated and plotted in **Supplementary Figure 8** for NAM-bound GPCRs and **Supplementary Figure 9** for PAM-bound GPCRs. Next, the residue contacts which exhibited the highest contact gradients (≥0.7) in the attention maps from DL and significant changes in fluctuations of the involved residues upon allosteric modulator binding were considered important for allosteric modulation of GPCRs (**Figures 2** and **3**).

**Figure 6.**
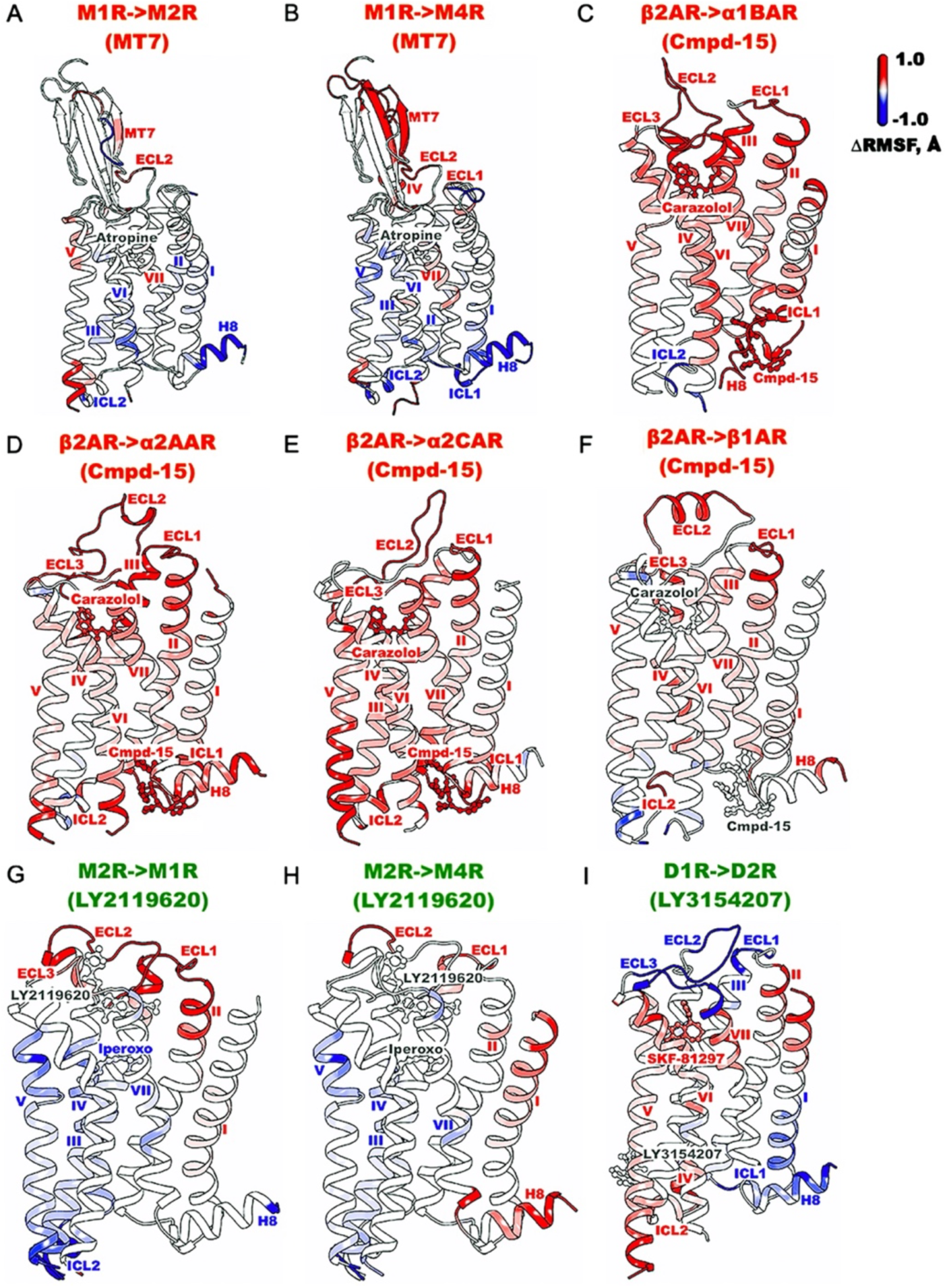
Increased system fluctuations were observed for binding of NAMs (MT7 and Cmpd-15) and PAMs (LY2119620 and LY3154207) to “non-cognate” GPCRs in GaMD simulations. Changes in root-mean-square fluctuations (ΔRMSFs) of the receptor, orthosteric and allosteric ligands in the M_2_R **(A)** and M_4_R **(B)** compared to the M_1_R bound by the MT7 NAM (PDB: 6WJC), the α_1B_AR **(C)**, α_2A_AR **(D)**, α_2C_AR **(E)**, and β_1_AR **(F)** compared to the β_2_AR bound by the Cmpd-15 NAM (PDB: 5X7D), the M_1_R **(G)** and M_4_R (PDB: 7V68) **(H)** compared to the M_2_R bound by the LY2119620 PAM (PDB: 6OIK), and the D_2_R **(I)** compared to the D_1_R bound by the LY3154207 PAM (PDB: 7LJC). A color scale of -1.0 (blue) to 0 (white) to 1.0 (red) is used to show the ΔRMSF.

The important residue contacts in NAM-bound GPCRs were mostly located near the allosteric binding sites (**Figure 2**). In the MT7-bound M_1_R, residue contacts V4.68-T172^ECL2^ and P5.36-T6.59 involved the TM4, TM5, and TM6 extracellular domains and ECL2 (**Figure 2A**). In the Cmpd-15-bound β_2_AR, residue contacts K6.29-P8.48 and T6.36-I7.52 involved the TM6 and TM7 intracellular domains and H8 (**Figure 2B**). In the AS408-bound β_2_AR, residue contact T4.56-V5.45 connected the middle of TM4 and TM5, while residue contact L34.56^ICL2^-I4.45 connected the TM4 intracellular domain and ICL2 (**Figure 2C**). In the NDT9513727-bound C5AR1, residue contact T3.45-A4.45 connected the TM3 and TM4 intracellular ends, while residue contact L5.51-F6.45 connected the middle of TM5 and TM6 (**Figure 2D**). In the Avacopan-bound C5AR1, residue contacts L4.56-L5.45 and P4.59-V5.38 connected the TM4 and TM5 extracellular ends (**Figure 2E**). In the ORG27569-bound CB_1_, residue contact R34.55^ICL2^-T4.38 connected the TM4 intracellular domain and ICL2, while residues T7.47-L7.55 involved the TM7 intracellular end (**Figure 2F**). In the GTPL9431-bound CCR2, residue contacts I5.61-K6.28 and V6.36-V7.56 connected the TM5, TM6, and TM7 intracellular domains (**Figure 2G**). In the NNC0640-bound GLP1R, residue contact L12.49^ICL1^-V8.50 connected ICL1 and H8, and residues E4.38-W4.40 involved the TM4 intracellular domain (**Figure 2H**). In the PF-06372222-bound GLP1R, residues Q210^ECL1^-H212^ECL1^ involved the ECL1, while residue contact L6.49-Q7.49 connected the middle of TM6 and TM7 (**Figure 2I**). Lastly, in the MK-0893-bound GLR, residue contact F5.51-I6.46 connected the middle of TM5 and TM6, while residue contact D6.61-R7.35 connected the extracellular domains of TM6 and TM7 (**Figure 2J**).

In PAM-bound GPCRs, important residue contacts were mostly located in the extracellular domains, orthosteric agonist-binding pocket, and intracellular G protein-binding regions (**Figure 3**). In the MIPS521-bound A_1_AR, residue contact G2.68-K168^ECL2^ involved the TM2 extracellular end and ECL2, while residue contact W6.48-L7.41 connected the middle of TM6 and TM7, interacting with the adenosine agonist (**Figure 3A**). In the LY2119620-bound M_2_R, residue contact M6.54-G7.38 were found interacting with the iperoxo agonist near the TM6 and TM7 extracellular ends, while residue contact C7.56-T8.49 connected the TM7 intracellular domain and H8 (**Figure 3B**). In the LY2119620-bound M_4_R, residue contacts N1.60-T8.53 and S5.62-T6.34 involved the TM1, TM5, and TM6 intracellular ends and H8 (**Figure 3C**). In the Cmpd-6FA-bound β_2_AR, residue contact T2.39-K4.39 connected the TM2 and TM4 intracellular ends, while residue contact K2.68-E7.33 connected the TM2 and TM7 extracellular domains (**Figure 3D**). In the LY3154207-bound D_1_R, residues V1.31-I1.43 and P34.50^ICL2^-K34.56^ICL2^ involved the TM1 extracellular end and ICL2, respectively (**Figure 3E**). In the AgoPAM-bound FFAR1, residue contact P5.32-N252^ECL3^ connected the TM5 extracellular domain and ECL3, while residue contact P34.50^ICL2^-F34.56^ICL2^ formed the short helix of ICL2 (**Figure 3F**). In the INT777-bound GPBAR, residues L4.59-G4.63 and residue contact W149^ECL2^-N154^ECL2^ involved the TM4 extracellular domain and ECL2, respectively (**Figure 3G**). In the LSN3160440-bound GLP1R, residue contacts A208^ECL1^-L217^ECL1^ and Q221^ECL1^-E294^ECL2^ connected ECL1 and ECL2 (**Figure 3H**).

### Free energy profiling of important residue contacts in GPCR allostery

2D free energy profiles were calculated from GaMD simulations for the important residue contacts identified from DL and structural flexibility analyses of GPCRs (**Figures 4** and **5**). Overall, binding of NAMs and PAMs reduced conformational space of the inactive antagonist-bound and active agonist-bound GPCRs, respectively. Moreover, in case the modulator-free GPCRs were able to sample multiple low-energy conformational states, binding of the NAMs and PAMs confined the GPCR residue contacts to fewer low-energy states (only 1 for most GPCRs) (**Figures 4** and **5**). The residue distances/RMSDs identified at the energy minima of these conformational states are listed in detail in **Supplementary Tables 3** and **4**.

Binding of the MT7 NAM to M_1_R confined the TM4 extracellular domain and ECL2 from two conformational states (“S1” and “S2”) to only the “S1” state, in which the extracellular mouth adopted the more closed conformation (**Figure 4A**). Cmpd-15 binding to the β_2_AR reduced the conformational space from three states (“S1”-”S3”) to only the “S3” state, in which the intracellular pocket formed by TM6, TM7, and H8 adopted the more open conformation to accommodate the NAM (**Figures 4B** and **2B**). Similarly, AS408 binding to the β_2_AR confined the TM4 intracellular end and ICL2 from two conformational states (“S1” and “S2”) to only state “S1”, in which the intracellular pocket formed by the TM4 intracellular end and ICL2 adopted a more open conformation for stable NAM binding (**Figures 4C** and **2C**). Binding of NDT9513727 to the C5AR1 reduced the number of low-energy conformational states from two to one (the “S1” state), where the allosteric pocket located between TM3 and TM4 intracellular ends as well as the middle of TM5 and TM6 adopted a more open conformation to accommodate the NAM (**Figures 4D** and **2D**). Avacopan-binding to the C5AR1 confined the TM4 and TM5 extracellular domains from two conformational states to only state “S1”, where the TM4 and TM5 extracellular ends adopted the more closed conformation to stabilize NAM binding (**Figures 4E** and **2E**). The hydrophobic stacking found between TM4 and TM5 extracellular ends was again in good agreement with previous finding by Xiaoli et al.^72^. Binding of MK-0893 to the GLR confined the conformational space from three states (“S1”-S3”) to only state “S1”, in which the middle of TM5 and TM6 as well as TM6 and TM7 extracellular ends adopted the more closed conformations (**Figure 4J**). In the cases of ORG27569-bound CB_1_, GTPL9431-bound CCR2, NNC0640-bound GLP1R, and PF-06372222-bound GLP1R, the modulator-free GPCRs already sampled only one low-energy conformational state (**Figure 4F-4I**).

In PAM-bound GPCRs, binding of MIPS521 to the A_1_AR confined the TM2 extracellular domain and ECL2 as well as the middle of TM6 and TM7 from three conformational states (“S1”-“S3”) to only the “S1” state, in which the extracellular mouth and orthosteric agonist-binding pocket adopted the more closed conformation to stabilize agonist binding (**Figures 5A** and **3A**). This finding was highly consistent with previous experimental and computational studies of the A_1_AR allosteric modulation^24,52,73,74^, where the PAM was found to significantly stabilize the adenosine agonist. The modulator-free GPCR in case of LY2119620-bound M_2_R sampled only one low-energy conformational state (**Figure 5B**). LY2119620-binding to the M_4_R reduced the conformational space of the TM1 intracellular end and H8 from three states (“S1”-”S3”) to only state “S1”, in which the G-protein-binding region adopted the more closed conformation to stabilize G-protein binding (**Figure 5C**). Binding of Cmpd-6FA to the β_2_AR confined the TM2 and TM7 extracellular ends as well as TM2 and TM4 intracellular ends from three states (“S1”-S3”) to the “S1” state, in which G protein-binding domain adopted a closed conformation (**Figure 5D**). Binding of the LY3154207 PAM to the D_1_R confined the TM1 extracellular end and ICL2 from two conformational states (“S1” and “S2”) to only state “S1”, reducing the flexibility of both domains (**Figures 5E** and **3E**). AgoPAM-binding to the FFAR1 reduced the conformational space of the TM5 extracellular end, ECL3, and ICL2 from two states (“S1” and “S2”) to state “S1”, where the extracellular mouth between TM5 and ECL3 as well as the ICL2 adopted a closed and short helix conformation, respectively (**Figures 5F** and **3F**). Here, the finding that the ICL2 preferred a short helix conformation in the AgoPAM-bound FFAR1 resembled the structural data of 5TZY and 5TZR PDB structures well^20^. Binding of INT777 to the allosteric site of GPBAR reduced the number of low-energy conformational states from three (“S1”-”S3”) to two (“S1” and “S2”) (**Figure 5G**). Lastly, LSN3160440-binding to the GLP1R confined ECL1 and ECL2 from two conformational states (“S1” and “S2”) to only the “S1” state, in which the extracellular mouth between ECL1 and ECL2 adopted the closed conformation to stabilize the peptide agonist (**Figures 5H** and **3H**).

### Selectivity of GPCR allosteric modulators

Additional GaMD simulations were performed on artificially generated computational models to examine binding selectivity of the MT7 and Cmpd-15 NAMs to the muscarinic and adrenergic receptors, respectively, as well as the LY2119620 and LY3154207 PAMs to the muscarinic and dopamine receptors, respectively. Flexibility changes were calculated by subtracting RMSFs of the cognate from the “non-cognate” GPCRs of the modulators (**Figure 6**). Furthermore, 2D free energy profiles of the heavy-atom RMSDs of orthosteric and allosteric ligands relative to their respective starting structures were calculated and shown in **Supplementary Figure 10**. Overall, modulator binding in the “non-cognate” GPCRs resulted in higher complex fluctuations and mostly larger conformational space compared to the cognate GPCRs, demonstrating the binding preference and selectivity of allosteric modulators towards their cognate subtypes.

Significantly higher fluctuations were observed for binding of allosteric modulators to “non-cognate” GPCRs, especially in the allosteric pockets and various receptor domains, compared to their binding to the cognate GPCRs (**Figure 6**). Compared to the MT7-bound M_1_R, the NAM showed moderately increased to much higher fluctuations in the model M_2_R and M_4_R, respectively. Furthermore, NAM binding increased fluctuations in the ECL2 of the M_2_R (**Figure 6A**) and the TM4 extracellular end, ECL1, and ECL2 in the M_4_R (**Figure 6B**). Compared to the Cmpd-15-bound β_2_AR, the NAM showed much higher fluctuations in the “non-cognate” subtypes of the α_1B_AR, α_2A_AR, and α_2C_AR, and significantly increased fluctuations of these three GPCR-antagonist complexes (**Figure 6C-6E**). The flexibility increase was smaller in the Cmpd-15-bound β_1_AR, likely due to the receptor similarity in its sequence and structure to the β_2_AR. Even so, binding of Cmpd-15 to the β_1_AR significantly increased fluctuations in the TM2 extracellular end, ECL1, ECL2, TM4, ICL2, and H8 (**Figure 6F**). Schober et al. uncovered that the binding preference of the LY2119620 PAM reduced from the M_2_R to the M_4_R and then M_1_R^75^. Here, LY2119620 binding to the M_1_R significantly increased fluctuations in the TM2, TM3 and TM4 extracellular ends, ECL1, ECL2, and ECL3 compared to the LY2119620-bound M_2_R (**Figure 6G**). In the LY2119620-bound M_4_R, PAM binding only slightly increased fluctuations in the TM2 and TM3 extracellular ends, ECL1, and ECL2 compared to the M_2_R (**Figure 6H**). Our simulation results were thus consistent with the experimental finding by Schober et al.^75^. Lastly, binding of the LY3154207 PAM to the D_2_R increased fluctuations mostly in the TM1 and TM2 extracellular ends, TM4 intracellular end, ICL2, TM5, TM6, TM7, and the SKF-81297 agonist compared to the cognate D_1_R (**Figure 6I**).

Binding of allosteric modulators to “non-cognate” GPCRs mostly increased conformational space of the orthosteric and allosteric ligands with higher RMSDs. Moreover, most of the modulator-bound “non-cognate” GPCRs sampled more low-energy conformational states compared to the cognate GPCRs (**Supplementary Figure 10**). In particular, the MT7-bound M_4_R sampled four low-energy conformational states (“S1”-”S4”) of the atropine antagonist and MT7 NAM compared to only two conformations (“S1” and “S2”) of the MT7-bound M_1_R (**Supplementary Figure 10A, 10C**). While the MT7-bound M_2_R sampled the same number of conformational states as the M_1_R, the “S3” state of the MT7-bound M_2_R showed higher RMSD of the atropine antagonist at ∼3.8 Å compared to the ∼1.0Å atropine RMSD in the “S1” state of M_1_R (**Supplementary Figure 10A-10B** and **Supplementary Table 5**). The Cmpd-15-bound α_1B_AR and α_2C_AR sampled three low-energy conformational states (“S2”-”S4”) compared to only one state (“S1”) of the Cmpd-15-bound β_2_AR (**Supplementary Figure 10D, 10E, 10G**). While the Cmpd-15-bound α_2A_AR sampled the same number of low-energy conformational states as the cognate β_2_AR, the “S3” state of the α_2A_AR showed higher RMSDs of both the carazolol antagonist and Cmpd-15 NAM at ∼2.3 Å and ∼6.0 Å, respectively, compared to the ∼1.0Å and ∼2.0Å carazolol and Cmpd-15 RMSDs of state “S1” of the β_2_AR (**Supplementary Figure 10D, 10F** and **Supplementary Table 5**). The LY2119620-bound M_1_R sampled only one low-energy conformational state (“S2”) compared to two states in the M_2_R (“S1” and “S2”) and M_4_R (“S3” and “S4”) (**Supplementary Figure 10I-10K**). Nevertheless, in the “S2” state, the LY2119620 PAM adopted the ∼6.0Å relatively higher RMSD conformation (**Supplementary Table 5**). Lastly, the LY3154207-bound D_2_R sampled two low-energy conformational states (“S2” and “S3”), both of which exhibited significantly higher agonist and PAM RMSDs, compared to the one “S1” state of LY3154207-bound D_1_R (**Supplementary Figure 10L-10M** and **Supplementary Table 5**).

## Conclusions

Allosteric modulators have emerged as more selective drug candidates than orthosteric agonist and antagonist ligands. However, many X-ray and cryo-EM structures of GPCRs resolved so far exhibit negligible differences upon binding of allosteric modulators. Consequently, mechanism of dynamic allosteric modulation in GPCRs remains unclear, despite their critical importance. In this work, we have integrated GaMD and DL in GLOW to map dynamic changes in free energy landscapes of GPCRs upon binding of allosteric modulators. By intersecting DL-predicted residue contacts with the highest gradient and residues with the largest flexibility changes, characteristic residue contacts were selected for free energy profiling to decipher the effects of allosteric modulator binding on GPCRs. The PAM and NAM binding primarily reduced fluctuations of the GPCR complexes. NAMs stabilized the allosteric and antagonist-binding sites. PAMs stabilized the receptor extracellular domains, orthosteric agonist-binding pocket, and G protein coupling regions. Furthermore, the conformational space of GPCRs was significantly reduced upon modulator binding. The NAMs and PAMs confined the GPCRs to mostly one specific conformation for signaling. These effects transcended across class A and B GPCRs. Furthermore, NAM and PAM binding were found selective towards their cognate receptor subtypes. Significantly higher fluctuations were observed for modulator binding to “non-cognate” GPCR subtypes, for which the orthosteric and allosteric ligands exhibited larger RMSDs.

While the effects of NAM and PAM binding on receptor dynamics were consistent across different GPCRs, exceptions were observed in the GPBAR bound by the INT777 PAM and the GLP1R bound by the PF-06372222 NAM. In the GPBAR, the fact that two molecules of the same charged ligand bound to the receptor could potentially create electrostatic repulsion, leading to increased fluctuations in the orthosteric ligand and other parts of the receptor^45^ (**Figure 3G**). In addition, potential inaccuracies in especially the ligand force field parameters could contribute to the inconsistencies observed in these two cases. Further ligand parameter optimization could be helpful to achieve more consistent results in these GPCR systems. In conclusion, we have deciphered the mechanism of dynamic allostery in class A and B GPCRs through DL of extensive GaMD simulations. Our findings are expected to facilitate rational design of selective GPCR allosteric drugs.

## Author Contributions

H.N.D. performed research, analyzed data, and wrote the manuscript. J.W. supervised GaMD simulations. Y.M. supervised the project, interpreted data, and wrote the manuscript. All authors contributed towards the final version of the manuscript.

## Acknowledgements

We thank Miao Lab members for valuable discussions. This work used supercomputing resources with allocation award TG-MCB180049 through the Advanced Cyberinfrastructure Coordination Ecosystem: Services & Support (ACCESS) program, which is supported by National Science Foundation grants #2138259, #2138286, #2138307, #2137603, and #2138296, and project M2874 through the National Energy Research Scientific Computing Center (NERSC), which is a U.S. Department of Energy Office of Science User Facility operated under Contract No. DE-AC02-05CH11231, and the Research Computing Cluster and BigJay Cluster funded through NSF Grant MRI-2117449 at the University of Kansas. This work was supported by the National Institutes of Health (R01GM132572).

