## Supplementary Information for "Deep Learning Dynamic Allostery of G-Protein-Coupled Receptors"

**Supplementary Figure 1. Time courses of the RMSDs of orthosteric ligands and NAMs relative to the respective starting structures calculated from GaMD simulations of class A GPCRs in the (A)** $M_1R$ without and with MT7, **(B)** $\beta_2AR$ without and with Cmpd-15, **(C)** $\beta_2AR$ without and with AS408, **(D)** C5AR1 without and with NDT9513727, **(E)** C5AR1 without and with Avacopan, **(F)** CB1 without and with ORG27569, and **(G)** CCR2 without and with GTPL9431. From left to right for each system are the RMSDs of the orthosteric ligand in the absence and presence of NAM (denoted "Antagonist" or "Agonist" and "AntagonistNAM" or "AgonistNAM", respectively) and RMSD of the NAM. The averages and standard deviations of the ligand RMSDs were included in each panel in the parentheses.

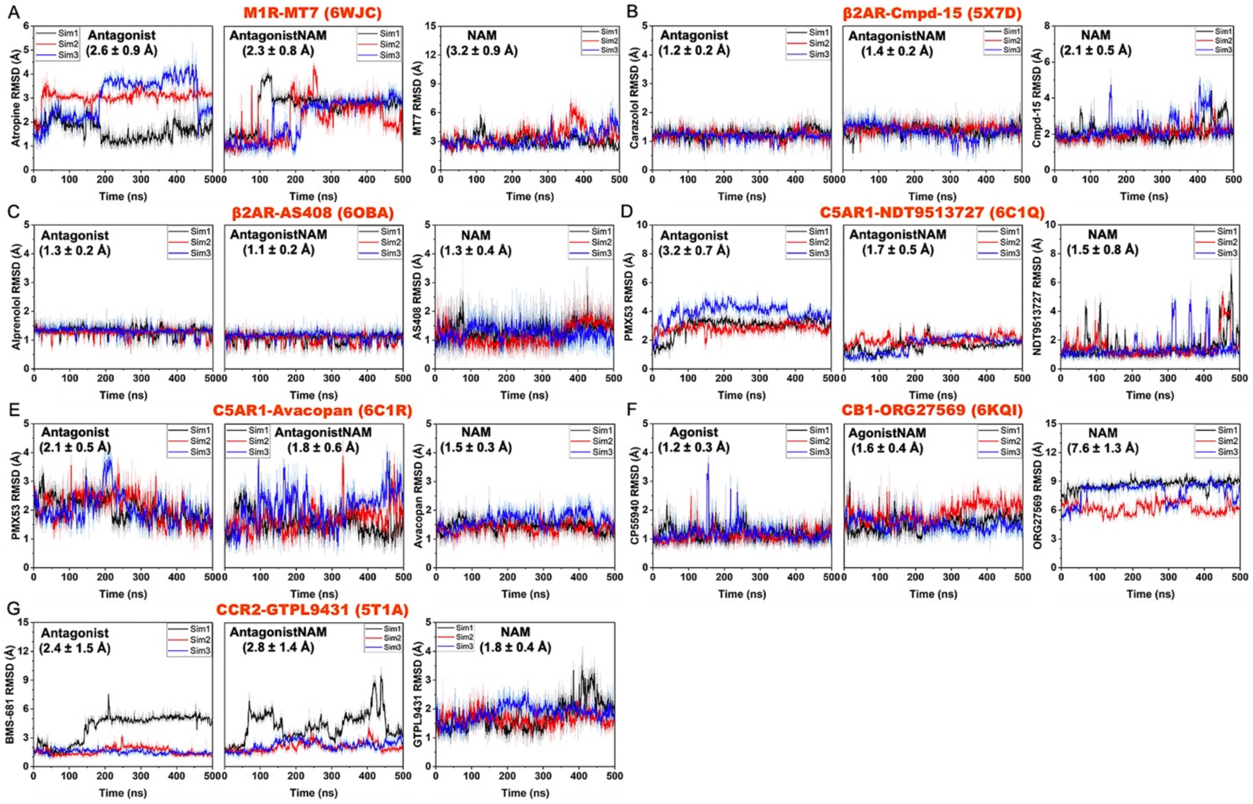

**Supplementary Figure 2. Time courses of the RMSDs of orthosteric ligands and PAMs relative to the respective starting structures calculated from GaMD simulations of class A GPCRs in the (A)** $A_1AR$ without and with MIPS521, **(B)** $M_2R$ without and with LY2119620, **(C)** $M_4R$ without and with LY2119620, **(D)** $\beta_2AR$ without and with Cmpd-6FA, **(E)** $D_1R$ without and with LY3154207, **(F)** FFAR1 without and with AgoPAM, and **(G)** GPBAR without and with INT777. From left to right for each system are the RMSDs of the orthosteric ligand in the absence and presence of PAM (denoted "Agonist" and "AgonistNAM", respectively) and RMSDs of the PAM. The averages and standard deviations of the ligand RMSDs were included in each panel in the parentheses.

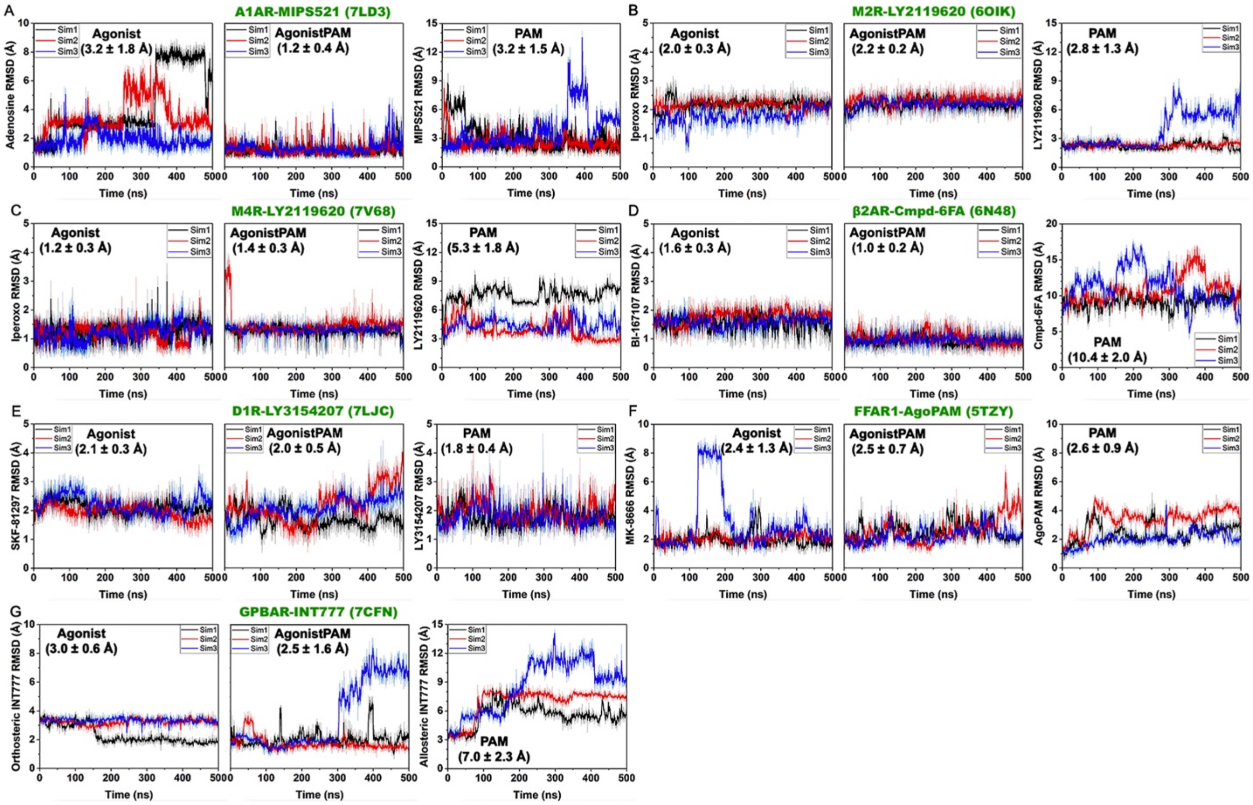

**Supplementary Figure 3. Time courses of the RMSDs of orthosteric and allosteric ligands relative to the respective starting structures calculated from GaMD simulations of class B GPCRs in the (A)** GLP1R with the NNC0640 NAM, **(B)** GLP1R with the PF-06372222 NAM, **(C)** GLP1R without and with the LSN3160440 PAM, and **(D)** GLR with the MK-0893 NAM. From left to right for **(C)** and **(D)** are the RMSDs of orthosteric ligands in the absence and presence of allosteric ligands (denoted “Agonist” or “Antagonist” and “AgonistPAM” or “AntagonistNAM”, respectively) and RMSDs of allosteric ligands. The averages and standard deviations of the ligand RMSDs were included in each panel in the parentheses.

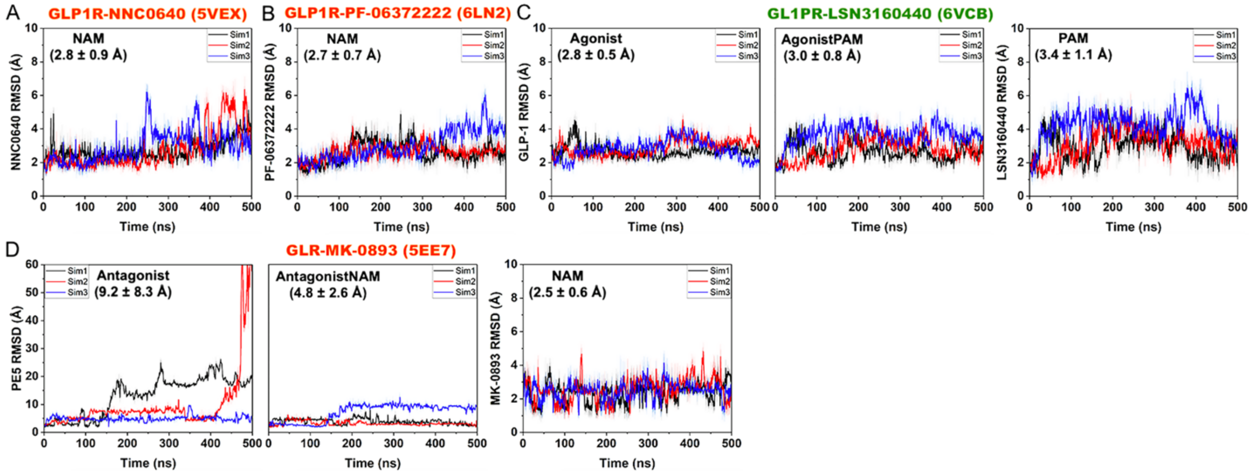

**Supplementary Figure 4. Changes in root-mean-square fluctuations (RMSFs) of receptors and orthosteric ligands of class A and B GPCRs upon binding of NAMs calculated from GaMD simulations** of the MT7-bound $M_1R$ (PDB: 6WJC) **(A)**, Cmpd-15-bound $\beta_2AR$ (PDB: 5X7D) **(B)**, AS408-bound $\beta_2AR$ (PDB: 6OBA) **(C)**, NDT9513727-bound C5AR1 (PDB: 6C1Q) **(D)**, Avacopan-bound C5AR1 (PDB: 6C1R) **(E)**, ORG27569-bound $CB_1$ (PDB: 6KQI) **(F)**, GTPL9431-bound CCR2 (PDB: 5T1A) **(G)**, NNC0640-bound GLP1R (PDB: 5VEX) **(H)**, PF-06372222-bound GLP1R (PDB: 6LN2) **(I)**, and MK-0893-bound GLR (PDB: 5EE7) **(J)**. A color scale of -1.0 (blue) to 0 (white) to 1.0 (red) is used to show the changes in RMSFs, and NAMs are colored orange.

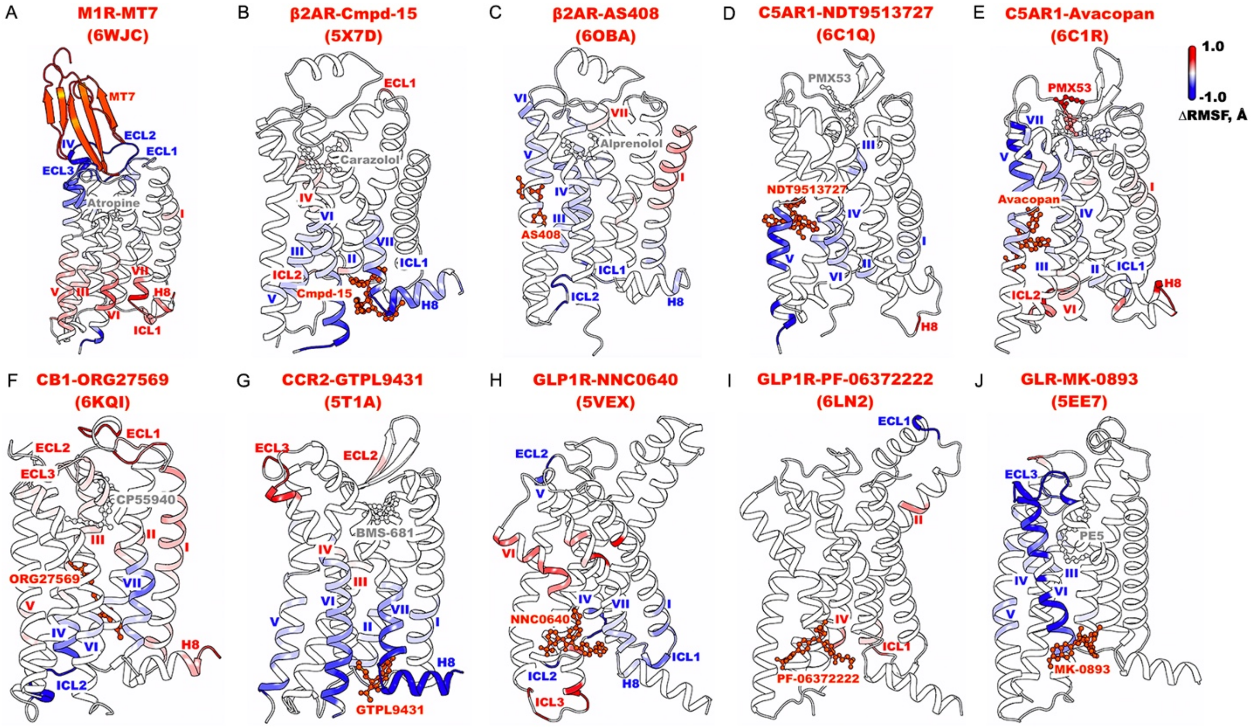

**Supplementary Figure 5. Changes in root-mean-square fluctuations (RMSFs) of receptors and orthosteric ligands of class A and B GPCRs upon binding of PAMs calculated from GaMD simulations** of the MIPS521-bound $A_1AR$ (PDB: 7LD3) **(A)**, LY2119620-bound $M_2R$ (PDB: 6OIK) **(B)**, LY2119620-bound $M_4R$ (PDB: 7V68) **(C)**, Cmpd-6FA-bound $\beta_2AR$ (PDB: 6N48) **(D)**, LY3154207-bound $D_1R$ (PDB: 7LJC) **(E)**, AgoPAM-bound FFAR1 (PDB: 5TZY) **(F)**, INT777-bound GPBAR (PDB: 7CFN) **(G)**, and LSN3160440-bound GLP1R **(H)**. A color scale of -1.0 (blue) to 0 (white) to 1.0 (red) is used to show the changes in RMSFs, and PAMs are colored green.

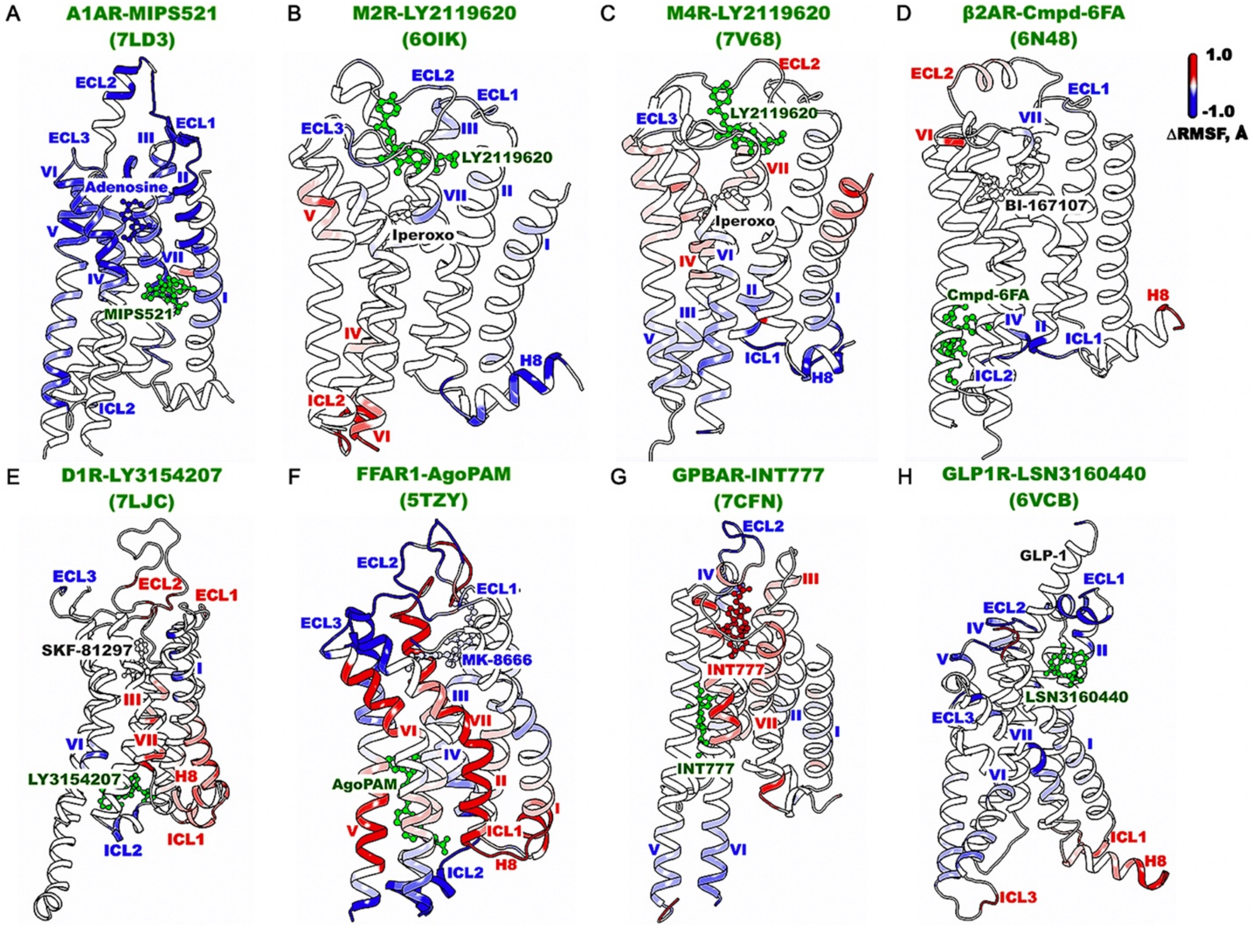

**Supplementary Figure 6. Accuracy curves of the training and validation datasets for GPCR allosteric modulation** of the $A_1AR$ **(A)**, muscarinic receptors $M_1R$, $M_2R$ and $M_4R$ **(B)**, $\beta_2AR$ **(C)**, C5AR1 **(D)**, $CB_1$ **(E)**, CCR2 **(F)**, $D_1R$ **(G)**, FFAR1 **(H)**, GPBAR **(I)**, GLP1R **(J)**, and GLR **(K)**.

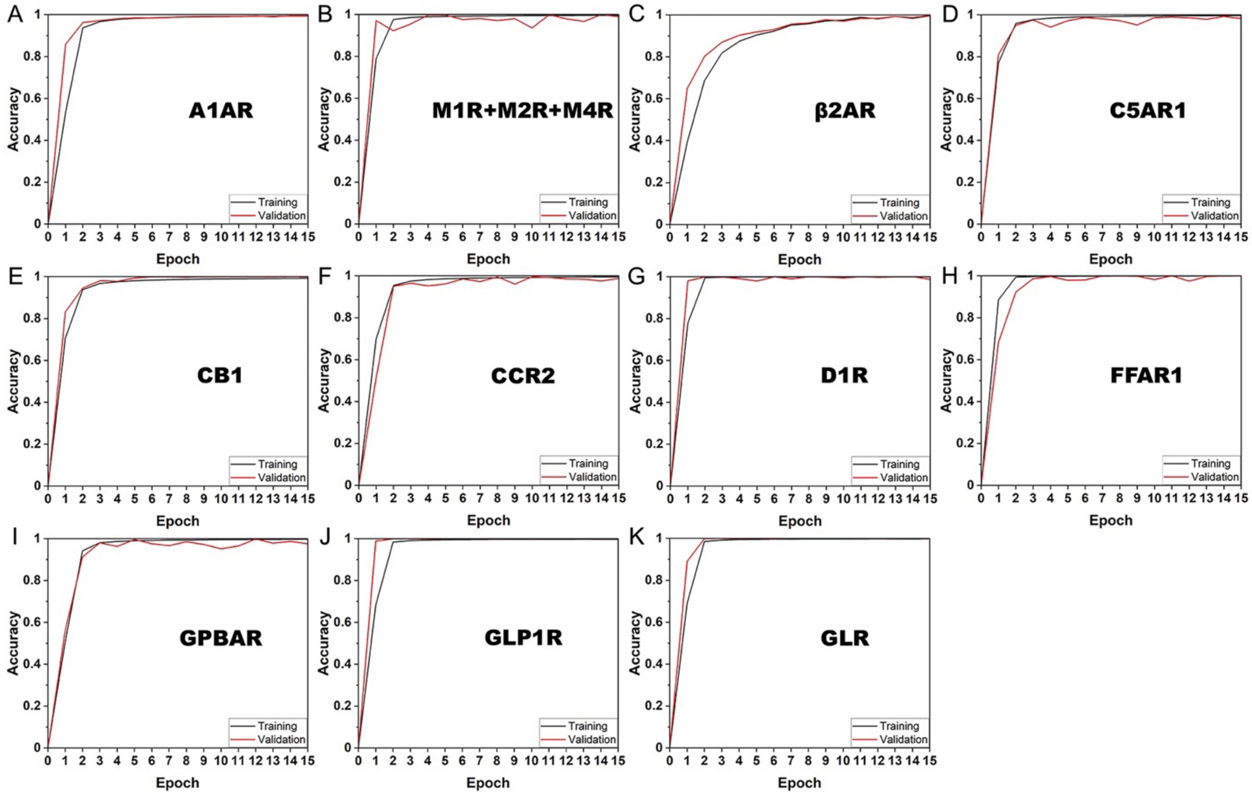

**Supplementary Figure 7. Confusion matrices calculated from the validation datasets for GPCR allosteric modulation** of the $A_1AR$ **(A)**, muscarinic receptors $M_1R$, $M_2R$ and $M_4R$ **(B)**, $\beta_2AR$ **(C)**, C5AR1 **(D)**, $CB_1$ **(E)**, CCR2 **(F)**, $D_1R$ **(G)**, FFAR1 **(H)**, GPBAR **(I)**, GLP1R **(J)**, and GLR **(K)**.

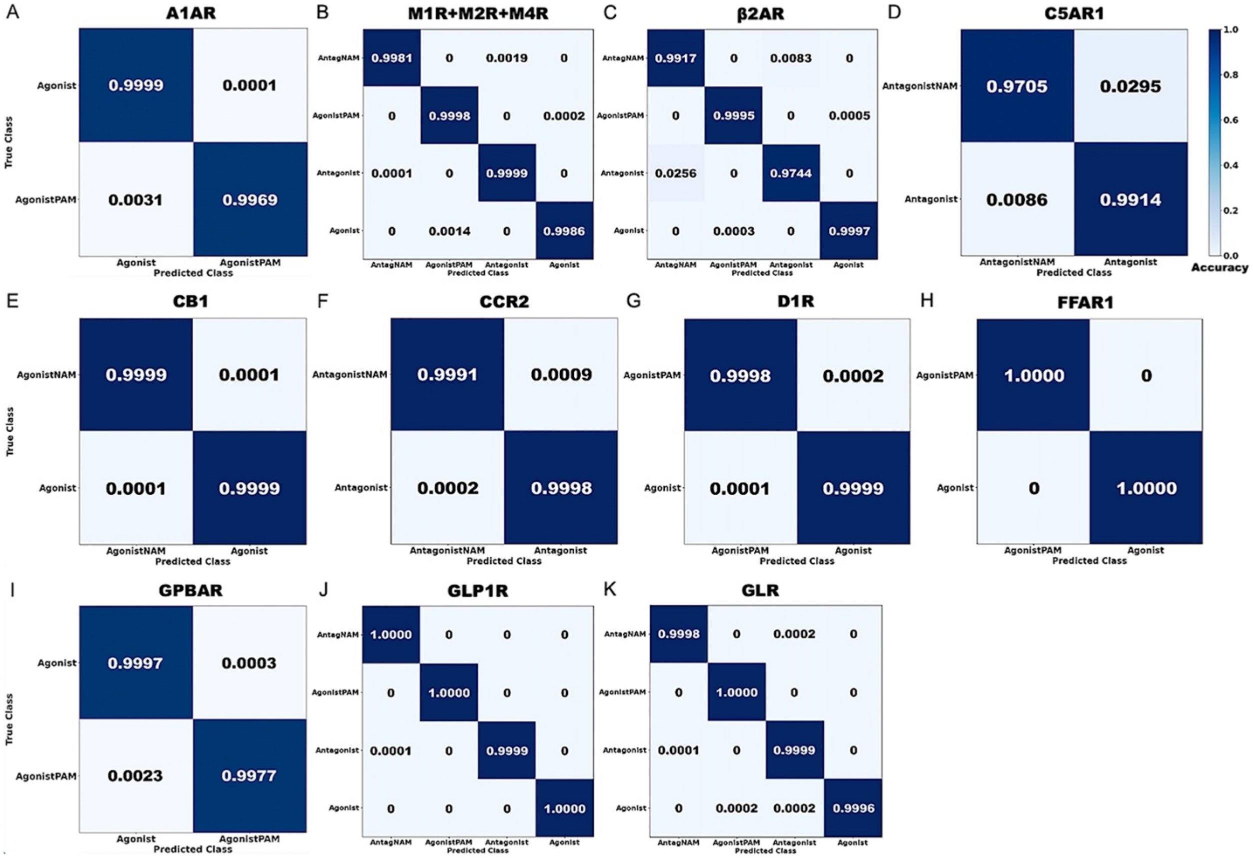

**Supplementary Figure 8. Saliency (attention) maps of residue contact gradients of class A and B GPCRs bound by negative allosteric modulators (NAMs),** including the MT7-bound $M_1R$ (PDB: 6WJC) **(A)**, Cmpd-15-bound $\beta_2AR$ (PDB: 5X7D) **(B)**, AS408-bound $\beta_2AR$ (PDB: 6OBA) **(C)**, NDT9513727-bound C5AR1 (PDB: 6C1Q) **(D)**, Avacopan-bound C5AR1 (PDB: 6C1R) **(E)**, ORG27569-bound $CB_1$ (PDB: 6KQI) **(F)**, GTPL9431-bound CCR2 (PDB: 5T1A) **(G)**, NNC0640-bound GLP1R (PDB: 5VEX) **(H)**, PF-06372222-bound GLP1R (PDB: 6LN2) **(I)**, and MK-0893-bound GLR (PDB: 5EE7) **(J)**. The seven transmembrane (TM) helices are labeled I-VII. Regions that contain residue contacts selected for free energy profiling are boxed in blue color. The gradients of residue contacts are shown in a 0.2 (white) to 0.25 (black) color scale.

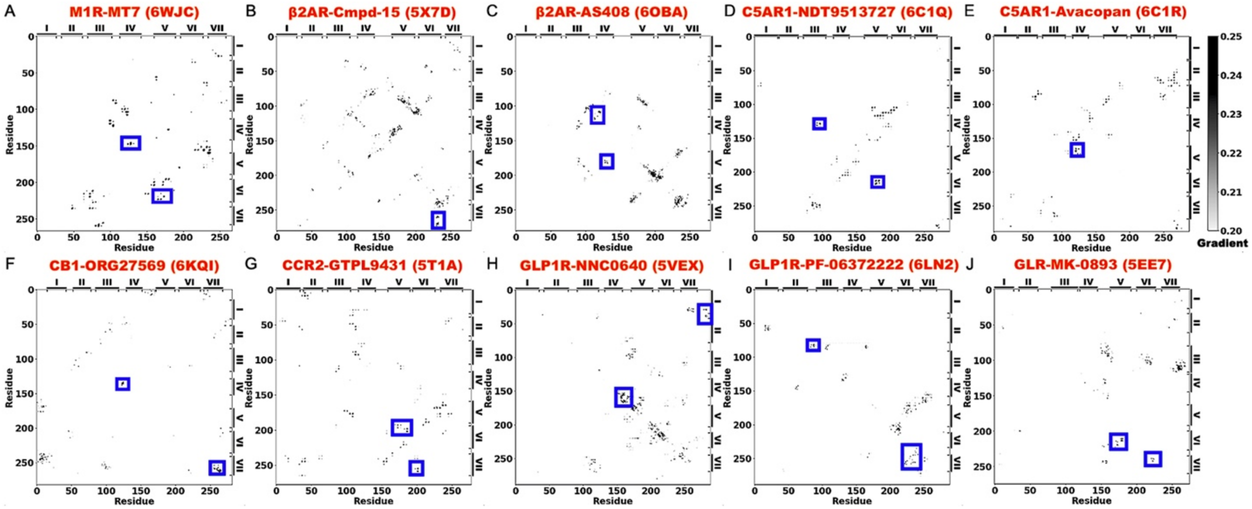

**Supplementary Figure 9. Saliency (attention) maps of residue contact gradients of class A and B GPCRs bound by positive allosteric modulators (PAMs),** including the MIPS521-bound $A_1$AR (PDB: 7LD3) **(A)**, LY2119620-bound $M_2$R (PDB: 6OIK) **(B)**, LY2119620-bound $M_4$R (PDB: 7V68) **(C)**, Cmpd-6FA-bound $\beta_2$AR (PDB: 6N48) **(D)**, LY3154207-bound $D_1$R (PDB: 7LJC) **(E)**, AgoPAM-bound FFAR1 (PDB: 5TZY) **(F)**, INT777-bound GPBAR (PDB: 7CFN) **(G)**, and LSN3160440-bound GLP1R **(H)**. The seven TM helices are labeled I-VII. Regions that contain residue contacts selected for free energy profiling are boxed in blue color. The gradients of residue contacts are shown in a 0.2 (white) to 0.25 (black) color scale.

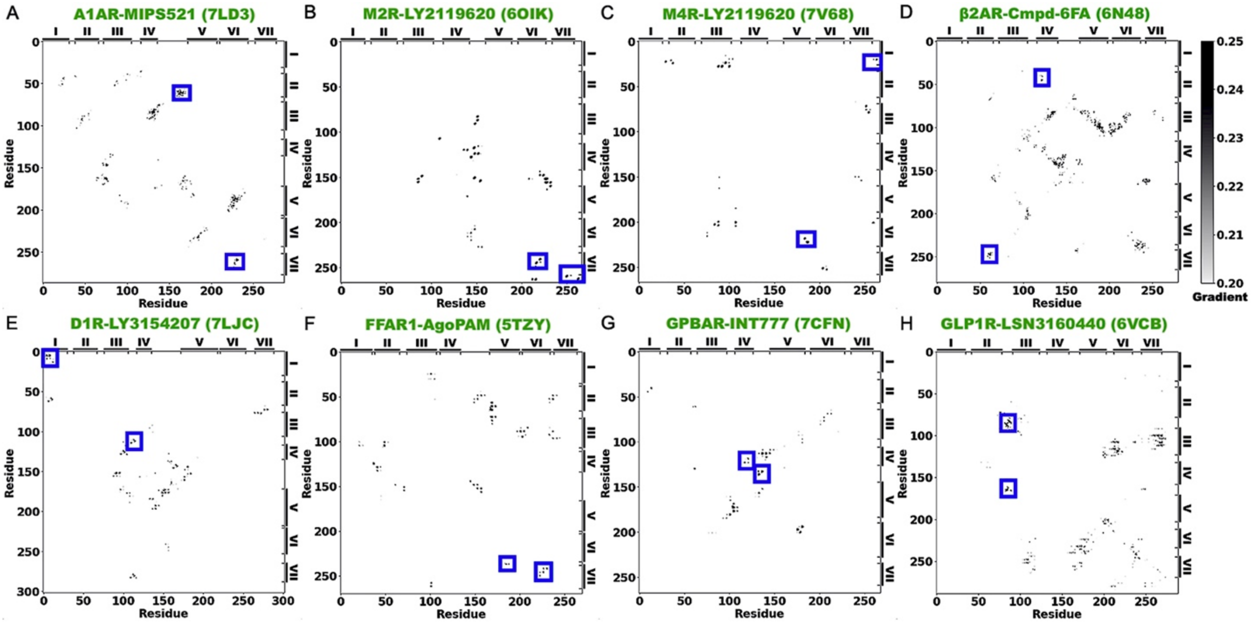

**Supplementary Figure 10. Reduced binding preference of NAMs (MT7 and Cmpd-15) and PAMs (LY2119620 and LY3154207) to "non-cognate" GPCRs. (A-C)** 2D free energy profiles of the heavy-atom RMSDs of the atropine antagonist and $C_\alpha$-atom RMSDs of the MT7 NAM relative to their starting structures in the $M_1R$ (PDB: 6WJC) **(A)**, model $M_2R$ **(B)**, and model $M_4R$ **(C)** bound by MT7. **(D-H)** 2D free energy profiles of the heavy-atom RMSDs of the carazolol antagonist and Cmpd-15 NAM relative to their starting structures in the $\beta_2AR$ (PDB: 5X7D) **(D),** model $\alpha_{1B}AR$ **(E)**, model $\alpha_{2A}AR$ **(F)**, model $\alpha_{2C}AR$ **(G)**, and model $\beta_1AR$ **(H)** bound by Cmpd-15. **(I-K)** 2D free energy profiles of the heavy-atom RMSDs of the iperoxo agonist and LY2119620 PAM relative to their starting structures in the $M_2R$ (PDB: 6OIK) **(I)**, $M_4R$ (PDB: 7V68) **(J)**, and model $M_1R$ **(K)** bound by LY2119620. **(L-M)** 2D free energy profiles of the heavy-atom RMSDs of the SKF-81297 agonist and LY3154207 PAM relative to their starting structures in the $D_1R$ (PDB: 7LJC) **(L)** and model $D_2R$ **(M)** bound by LY3154207.

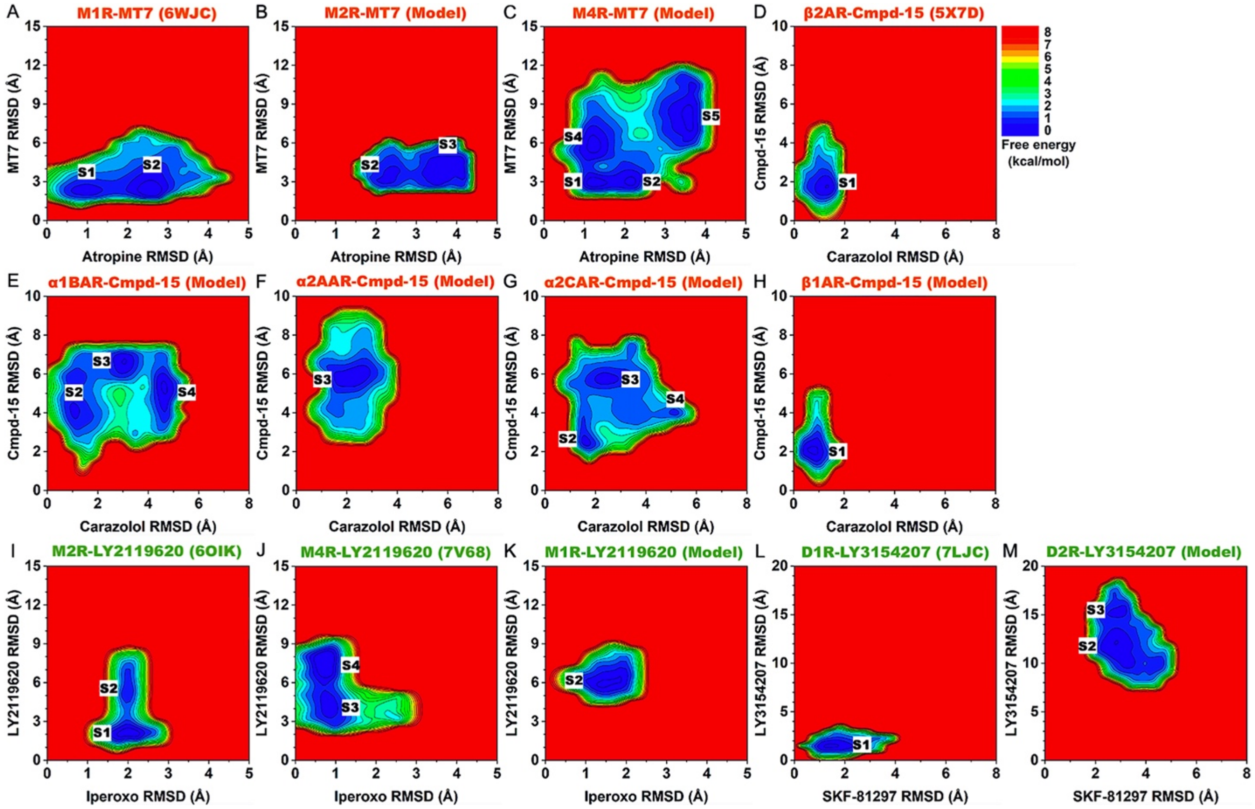

**Supplementary Table 1.** Summary of GaMD simulations performed on the PDB structures and computational models ("Model") of GPCRs with and without the positive/negative allosteric modulator (PAM/NAM). **(a)** PDB IDs or computational models ("Model") built for GaMD simulations. **(b)** Types of modulators bound in GPCRs. **(c)** Names and overall charges of orthosteric ligands bound in GPCRs. **(d)** Names and overall charges of allosteric ligands bound in GPCRs. **(e)** Averages and standard deviations of boost potentials $\Delta V$ (kcal/mol) calculated from three independent 500ns GaMD simulations for each GPCR system.

| Receptor | PDB/Model[a] | PAM/NAM[b] | Orthosteric ligand (charge)[c] | Allosteric ligand (charge)[d] | $\Delta V$ (kcal/mol) (3 x 500ns)[e] |
|---|---|---|---|---|---|
| $A_1$AR | 7LD3 | PAM | Adenosine (0) | MIPS521 (0) | 20.3 ± 7.2 |
| | | | | - | 19.0 ± 8.0 |
| $M_1$R | 6WJC | NAM | Atropine (0) | MT7 (0) | 13.1 ± 4.1 |
| | | | | - | 14.3 ± 4.3 |
| | Model | PAM | Iperoxo (+1) | LY2119620 (0) | 15.4 ± 4.5 |
| $M_2$R | Model | NAM | Atropine (0) | MT7 (0) | 14.0 ± 4.2 |
| | 6OIK | PAM | Iperoxo (+1) | LY2119620 (0) | 14.0 ± 4.3 |
| | | | | - | 15.0 ± 4.5 |
| $M_4$R | Model | NAM | Atropine (0) | MT7 (0) | 13.8 ± 4.2 |
| | 7V68 | PAM | Iperoxo (+1) | LY2119620 (0) | 14.6 ± 4.4 |
| | | | | - | 15.3 ± 4.8 |
| $\alpha_{1B}$AR | Model | NAM | Carazolol (+1) | Cmpd-15 (0) | 13.9 ± 4.2 |
| $\alpha_{2A}$AR | Model | NAM | Carazolol (+1) | Cmpd-15 (0) | 14.0 ± 4.3 |
| $\alpha_{2C}$AR | Model | NAM | Carazolol (+1) | Cmpd-15 (0) | 14.4 ± 4.3 |
| $\beta_1$AR | Model | NAM | Carazolol (+1) | Cmpd-15 (0) | 14.4 ± 4.3 |
| $\beta_2$AR | 5X7D | NAM | Carazolol (+1) | Cmpd-15 (0) | 14.5 ± 4.3 |
| | | | | - | 14.3 ± 4.3 |
| | 6OBA | NAM | Alprenolol (+1) | AS408 (+1) | 14.4 ± 4.3 |
| | | | | - | 14.7 ± 4.4 |
| | 6N48 | PAM | BI-167107 (+1) | Cmpd-6FA (-1) | 14.3 ± 4.3 |
| | | | | - | 14.5 ± 4.3 |
| C5AR1 | 6C1Q | NAM | PMX53 (0) | NDT9513727 (0) | 13.8 ± 4.2 |
| | | | | - | 14.2 ± 4.3 |
| | 6C1R | NAM | PMX53 (0) | Avacopan (0) | 13.2 ± 4.1 |
| | | | | - | 14.4 ± 4.3 |
| $CB_1$ | 6KQI | NAM | CP55940 (0) | ORG27569 (0) | 13.0 ± 4.1 |
| | | | | - | 13.6 ± 4.2 |
| CCR2 | 5T1A | NAM | BMS-681 (0) | GTPL9431 (0) | 13.9 ± 4.2 |
| | | | | - | 13.6 ± 4.2 |
| $D_1$R | 7LJC | PAM | SKF-81297 (+1) | LY3154207 (0) | 13.3 ± 4.2 |
| | | | | - | 14.6 ± 4.4 |
| $D_2$R | Model | PAM | SKF-81297 (+1) | LY3154207 (0) | 15.2 ± 4.5 |
| FFAR1 | 5TZY | PAM | MK-8666 (-1) | AgoPAM (-1) | 13.8 ± 4.2 |
| | | | | - | 14.1 ± 4.3 |
| GPBAR | 7CFN | PAM | INT777 (-1) | INT777 (-1) | 14.3 ± 4.3 |
| | | | | - | 15.3 ± 4.5 |
| GLP1R | 5VEX | NAM | - | NNC0640 (0) | 14.0 ± 4.3 |
| | | | | - | 14.4 ± 4.4 |
| | 6LN2 | NAM | - | PF-06372222 (-1) | 14.2 ± 4.3 |
| | | | | - | 14.5 ± 4.3 |
| | 6VCB | PAM | GLP-1 (0) | LSN3160440 (+1) | 13.4 ± 4.2 |
| | | | | - | 15.8 ± 4.6 |
| GLR | 5EE7 | NAM | PE5 (0) | MK-0893 (-1) | 13.7 ± 4.3 |
| | | | | - | 15.0 ± 4.4 |

**Supplementary Table 2.** Characteristic residue contacts with ≥ 0.7 gradients identified from GLOW analysis of GPCR allosteric modulation. Residue contacts selected for free energy profiling are in bold.

| System | PDB/Model | PAM/NAM | Residue contacts |
|---|---|---|---|
| A$_1$AR-MIPS521 | 7LD3 | PAM | T23.52$^{ECL1}$ – P165$^{ECL2}$, N2.65 – C45.50$^{ECL2}$, C3.25 – K168$^{ECL2}$, Q23.51$^{ECL1}$ – C45.50$^{ECL2}$, Y5.58 – F6.42, L5.53 – S6.47, **G2.68 – K168$^{ECL2}$,** E5.36 – L6.59, **W6.48 – L7.41,** N148$^{ECL2}$ – V152$^{ECL2}$ |
| M$_1$R-MT7 | 6WJC | NAM | **V4.68 – T172$^{ECL2}$, P5.36 – T6.59**, W4.64 – Q177$^{ECL2}$ |
| M$_2$R-LY2119620 | 6OIK | PAM | G3.21 – D173$^{ECL2}$, P34.50$^{ICL2}$ – K34.56$^{ICL2}$, V6.33 – N8.47, V168$^{ECL2}$ – D173$^{ECL2}$, **M6.54 – G7.38,** P3.22 – D173$^{ECL2}$, **C7.56 – T8.49,** T8.53 – L8.57 |
| M$_4$R-LY2119620 | 7V68 | PAM | **N1.60 – T8.53, S5.62 – T6.34,** N1.60 – L12.50$^{ICL1}$, N1.60 – T2.37, P5.36 – T6.59, I6.40 – Y7.53 |
| $\beta_2$AR-Cmpd-15 | 5X7D | NAM | K34.52$^{ICL2}$ – K6.29, D3.49 – K6.29, R12.49$^{ICL1}$ – D8.49, E6.30 – R7.55, **T6.36 – I7.52, K6.29 – P8.48** |
| $\beta_2$AR-AS408 | 6OBA | NAM | S34.55$^{ICL2}$ – T4.38, Y34.53$^{ICL2}$ – L34.56$^{ICL2}$, **T4.56 – V5.45, L34.56$^{ICL2}$ – I4.45,** V3.33 – F4.58, E3.41 – V5.46, C6.27 – A6.33, E3.41 – S4.53 |
| $\beta_2$AR-Cmpd-6FA | 6N48 | PAM | **T2.39 – K4.39**, H178$^{ECL2}$ – N183$^{ECL2}$, **K2.68 – E7.33**, R3.50 – L6.34, A3.38 – W6.48, F45.52$^{ECL2}$ – L302$^{ECL3}$, R3.50 – F5.62, L34.56$^{ICL2}$ – I4.45 |
| C5AR1-NDT9513727 | 6C1Q | NAM | N34.55$^{ICL2}$ – S5.67, F3.55 – G4.40, N3.35 – N7.45, **T3.45 – A4.45, L5.51 – F6.45**, S3.26 – V7.39 |
| C5AR1-Avacopan | 6C1R | NAM | I2.59 – D7.35, Y4.63 – K5.32, R4.64 – R5.42, L3.43 – I7.51, **L4.56 – L5.45, P4.59 – V5.38** |
| CB$_1$-ORG27569 | 6KQI | NAM | F102 – V7.34, V110 – D266$^{ECL2}$, M103 – D266$^{ECL2}$, V110 – T7.33, L12.50$^{ICL1}$ – L8.50, **R34.55$^{ICL2}$ – T4.38,** I6.40 – I7.52, **T7.47 – L7.55** |
| CCR2-GTPL9431 | 5T1A | NAM | V6.43 – N7.49, **I5.61 – K6.28, V6.36 – V7.56**, W5.34 – F6.64, P5.50 – W6.48, I3.54 – R5.63, W5.34 – N276$^{ECL3}$, L34.54$^{ICL2}$ – R34.57$^{ICL2}$, R34.57$^{ICL2}$ – S5.62 |
| D$_1$R-LY3154207 | 7LJC | PAM | S172$^{ECL2}$ – E181$^{ECL2}$, N3.26 – S45.52$^{ECL2}$, K167$^{ECL2}$ – R5.36, P34.50$^{ICL2}$ – F7.55, V1.31 – E2.65, **P34.50$^{ICL2}$ – K34.56$^{ICL2}$, V1.31 – I1.43**, W163$^{ECL2}$ – N185$^{ECL2}$ |
| FFAR1-AgoPAM | 5TZY | PAM | **P5.32 – N252$^{ECL3}$, P34.50$^{ICL2}$ – F34.56$^{ICL2}$,** S6.54 – W7.34, G6.39 – V7.52, S154$^{ECL2}$ – T162$^{ECL2}$, R3.50 – T7.53, L3.43 – S7.45, G70$^{ECL1}$ – L45.51$^{ECL2}$ |
| GPBAR-INT777 | 7CFN | PAM | I1.47 – L2.51, G4.63 – G152$^{ECL2}$, W149$^{ECL2}$ – G152$^{ECL2}$, G152$^{ECL2}$ – A159$^{ECL2}$, **W149$^{ECL2}$ – N154$^{ECL2}$**, L5.69 – T6.30, I1.47 – L2.51, **L4.59 – G4.63,** P151$^{ECL2}$ – A5.36 |
| GLP1R-NNC0640 | 5VEX | NAM | I1.60 – C7.58, **L12.49$^{ICL1}$ – V8.50, E4.38 – W4.40,** N5.32 – N5.34, N300$^{ECL2}$ – W5.36, Q210$^{ECL1}$ – W45.51$^{ECL2}$, L6.48 – T7.46, L6.38 – L7.56 |
| GLP1R-PF-06372222 | 6LN2 | NAM | S219$^{ECL1}$ – L3.31, Y220$^{ECL1}$ – C45.50$^{ECL2}$, Y1.40 – D2.68, T2.45 – Q4.39, **Q210$^{ECL1}$ – H212$^{ECL1}$, L6.49 – Q7.49** |
| GLP1R-LSN3160440 | 6VCB | PAM | F5.54 – I6.46, **A208$^{ECL1}$ – L217$^{ECL1}$, Q221$^{ECL1}$ – E294$^{ECL2}$**, Y3.45 – F7.40, R5.40 – I6.46, L3.47 – Y7.57 |
| GLR-MK-0893 | 5EE7 | NAM | K4.64 – N291$^{ECL2}$, N298$^{ECL2}$ – K7.38, G3.39 – K7.38, V3.34 – W5.36, **F5.51 – I6.46, D6.61 – R7.35** |

**Supplementary Table 3. Summary of the 2D free energy profiles of characteristic residues in the allosteric modulation of class A and B GPCRs bound by NAMs in Figure 4.** Locations of the energy minima are included in the third and last columns, with coordinates of the first and second reaction coordinates listed.

| System | Reaction coordinates | Without NAM | With NAM |
|---|---|---|---|
| $M_1$R-MT7 | V4.68-T172 Distance, P5.36-T6.59 Distance | **S1**(~9.0Å, ~4.5Å) **S2**(~11.9Å, ~4.6Å) | **S1**(~9.5Å, ~5Å) |
| $\beta_2$AR-Cmpd-15 | T6.36-I7.52 Distance, K6.29-P8.48 Distance | **S1**(~6.7Å, ~6.5Å) **S2**(~6.7Å, ~10.0Å) **S3**(~7.5Å, ~11.0Å) | **S3**(~7.0Å, ~10.0Å) |
| $\beta_2$AR-AS408 | T4.56-V5.45 Distance, $L34.56^{ICL2}$-I4.45 Distance | **S1**(~8.0Å, ~12.5Å) **S2**(~7.2Å, ~8.0Å) | **S1**(~8.0Å, ~12.5Å) |
| C5AR1-NDT9513727 | T3.45-A4.45 Distance, L5.51-F6.45 Distance | **S1**(~7.0Å, ~7.0Å) **S2**(~7.0Å, ~10.5Å) | **S1**(~7.5Å, ~7.5Å) |
| C5AR1-Avacopan | L4.56-L5.45 Distance, P4.59-V5.38 Distance | **S1**(~9.0Å, ~7.0Å) **S2**(~11.8Å, ~11.0Å) | **S1**(~8.5Å, ~6.5Å) |
| CB1-ORG27569 | $R34.55^{ICL2}$-T4.38 Distance, T7.47-L7.55 RMSD | **S1**(~6.0Å, ~0.5Å) | **S1**(~6.0Å, ~0.5Å) |
| CCR2-GTPL9431 | I5.61-K6.28 Distance, V6.36-V7.56 Distance | **S1**(~7.5Å, ~6.5Å) | **S1**(~5.0Å, ~6.3Å) |
| GLP1R-NNC0640 | $L12.49^{ICL1}$-V8.50 Distance, E4.38-W4.40 RMSD | **S1**(~7.5Å, ~2.0Å) | **S1**(~7.5Å, ~1.5Å) |
| GLP1R-PF-06372222 | $Q210^{ECL1}$-$H212^{ECL1}$ RMSD, L6.49-Q7.49 Distance | **S1**(~5.0Å, ~6.5Å) | **S1**(~2.5Å, ~7.5Å) |
| GLR-MK-0893 | F5.51-I6.46 Distance, D6.61-R7.35 Distance | **S1**(~7.0Å, ~9.0Å) **S2**(~7.5Å, ~12.0Å) **S3**(~11.9Å, ~8.5Å) | **S1**(~7.5Å, ~9.5Å) |

**Supplementary Table 4. Summary of the 2D free energy profiles of characteristic residues in the allosteric modulation of class A and B GPCRs bound by PAMs in Figure 5.** Locations of the energy minima are included in the third and last columns, with coordinates of the first and second reaction coordinates listed.

| System | Reaction coordinates | Without PAM | With PAM |
|---|---|---|---|
| $A_1$AR-MIPS521 | G2.68-K168$^{ECL2}$ Distance,<br>W6.48-L7.41 Distance | **S1**(~8.0Å, ~7.5Å)<br>**S2**(~5.5Å, ~10.0Å)<br>**S3**(~11.5Å, ~10.0Å) | **S1**(~5.5Å, ~7.0Å) |
| $M_2$R-LY2119620 | M6.54-G7.38 Distance,<br>C7.56-T8.49 Distance | **S1**(~8.0Å, ~9.0Å) | **S1**(~8.0Å, ~9.0Å) |
| $M_4$R-LY2119620 | N1.60-T8.53 Distance,<br>S5.62-T6.34 Distance | **S1**(~10.0Å, ~5.5Å)<br>**S2**(~6.0Å, ~5.5Å)<br>**S3**(~11.0Å, ~8.0Å) | **S1**(~7.0Å, ~5.5Å) |
| $\beta_2$AR-Cmpd-6FA | T2.39-K4.39 Distance,<br>K6.28-E7.33 Distance | **S1**(~9.0Å, ~12.0Å)<br>**S2**(~15.5Å, ~11.5Å)<br>**S3**(~12.5Å, ~7.5Å) | **S1**(~9.5Å, ~12.0Å) |
| $D_1$R-LY3154207 | V1.31-I1.43 RMSD,<br>P34.50$^{ICL2}$-K34.56$^{ICL2}$ RMSD | **S1**(~2.0Å, ~2.5Å)<br>**S2**(~5.0Å, ~2.0Å) | **S1**(~2.0Å, ~1.0Å) |
| FFAR1-AgoPAM | P5.32-N252 Distance,<br>P34.50$^{ICL2}$-F34.56$^{ICL2}$ Distance | **S1**(~7.0Å, ~9.5Å)<br>**S2**(~14.0Å, ~13.0Å)<br>**S3**(~13.0Å, ~8.5Å) | **S1**(~7.0Å, ~9.5Å) |
| GPBAR-INT777 | L4.59-G4.63 RMSD,<br>W149$^{ECL2}$-N154$^{ECL2}$ Distance | **S1**(~2.0Å, ~3.0Å)<br>**S2**(~3.0Å, ~10.0Å)<br>**S3**(~7.0Å, ~5.0Å) | **S1**(~2.0Å, ~3.5Å)<br>**S2**(~3.0Å, ~8.0Å) |
| GLP1R-LSN3160440 | A208$^{ECL1}$-L217$^{ECL1}$ Distance,<br>Q221$^{ECL1}$ – E294$^{ECL2}$ Distance | **S1**(~5.5Å, ~7.5Å)<br>**S2**(~7.5Å, ~8.0Å) | **S1**(~5.5Å, ~7.5Å) |

**Supplementary Table 5. Summary of the 2D free energy profiles of orthosteric and allosteric ligand binding in the $M_1R$, $M_2R$, and $M_4R$ bound by the MT7 NAM, $\beta_2AR$, $\alpha_{1B}AR$, $\alpha_{2A}AR$, $\alpha_{2C}AR$, and $\beta_1AR$ bound by the Cmpd-15 NAM, $M_2R$, $M_4R$, and $M_1R$ bound by the LY2119620 PAM, and $D_1R$ and $D_2R$ bound by the LY3154207 PAM in Figure S10.** Locations of the energy minima are included in the third and last columns, with coordinates of the first and second reaction coordinates listed.

| System | Reaction coordinates | Low-energy states |
|---|---|---|
| $M_1R$-MT7 | Atropine RMSD,<br>MT7 RMSD | **S1**(~1.0Å, ~2.5Å)<br>**S2**(~2.5Å, ~2.5Å) |
| $M_2R$-MT7 | | **S2**(~2.3Å, ~3.5Å)<br>**S3**(~3.8Å, ~4.0Å) |
| $M_4R$-MT7 | | **S1**(~1.3Å, ~3.0Å)<br>**S2**(~2.3Å, ~3.0Å)<br>**S4**(~1.3Å, ~6.0Å)<br>**S5**(~3.5Å, ~8.0Å) |
| $\beta_2AR$-Cmpd-15 | Carazolol RMSD,<br>Cmpd-15 RMSD | **S1**(~1.0Å, ~2.0Å) |
| $\alpha_{1B}AR$-Cmpd-15 | | **S2**(~1.0Å, ~4.0Å)<br>**S3**(~3.0Å, ~6.5Å)<br>**S4**(~4.5Å, ~5.0Å) |
| $\alpha_{2A}AR$-Cmpd-15 | | **S3**(~2.3Å, ~6.0Å) |
| $\alpha_{2C}AR$-Cmpd-15 | | **S2**(~1.7Å, ~2.5Å)<br>**S3**(~2.3Å, ~5.7Å)<br>**S4**(~5.1Å, ~4.0Å) |
| $\beta_1AR$-Cmpd-15 | | **S1**(~0.8Å, ~2.0Å) |
| $M_2R$-LY2119620 | Iperoxo RMSD,<br>LY2119620 RMSD | **S1**(~2.0Å, ~2.0Å)<br>**S2**(~2.0Å, ~5.5Å) |
| $M_4R$-LY2119620 | | **S3**(~0.8Å, ~4.0Å)<br>**S4**(~0.8Å, ~7.5Å) |
| $M_1R$-LY2119620 | | **S2**(~1.5Å, ~6.0Å) |
| $D_1R$-LY3154207 | SKF-81297 RMSD,<br>LY3154207 RMSD | **S1**(~1.5Å, ~1.5Å) |
| $D_2R$-LY3154207 | | **S2**(~3.0Å, ~15.5Å)<br>**S3**(~3.0Å, ~11.5Å) |
